# HBO1-MLL interaction promotes AF4/ENL/P-TEFb-mediated leukemogenesis

**DOI:** 10.1101/2021.01.08.425834

**Authors:** Satoshi Takahashi, Akinori Kanai, Hiroshi Okuda, Ryo Miyamoto, Takeshi Kawamura, Hirotaka Matsui, Toshiya Inaba, Akifumi Takaori-Kondo, Akihiko Yokoyama

## Abstract

Leukemic oncoproteins cause uncontrolled self-renewal of hematopoietic progenitors by aberrant gene activation, eventually causing leukemia. However, the molecular mechanism of aberrant gene activation remains elusive. Here, we showed that leukemic MLL fusion proteins associate with the HBO1 histone acetyltransferase (HAT) complex through their TRX2 domain. Among many MLL fusions, MLL-ELL particularly depended on its association with the HBO1 complex for leukemic transformation. The C-terminal portion of ELL provided a binding platform for multiple factors including AF4, EAF1 and p53. MLL-ELL activated gene expression by loading an AF4 /ENL/P-TEFb complex (AEP) onto the target promoters. The HBO1 complex promoted the use of AEP over EAF1 and p53. Moreover, the NUP98-HBO1 fusion protein exerted its oncogenic properties via interaction with MLL but not its intrinsic HAT activity. Thus, the interaction between HBO1 and MLL is an important nexus in leukemic transformation, which may serve as a therapeutic target for drug development.

## Introduction

Mutated transcriptional regulators often cause uncontrolled self-renewal of immature hematopoietic precursors, which leads to aggressive leukemia. MLL (also known as KMT2A and MLL1) is a transcriptional maintenance factor that upregulates homeobox (*HOX*) genes in development (Yu *et al*, 1998). Chromosomal translocations of the *MLL* gene generate MLL fusion genes with more than 80 different partners to induce leukemia (Meyer *et al*, 2018). MLL fusion proteins cause uncontrolled self-renewal by constitutively activating various oncogenic genes (e.g., *HOXA9*, *MEIS1*), whose expression is normally restricted in immature precursors such as hematopoietic stem cells (HSCs) (Krivtsov *et al*, 2006). However, the mechanisms by which MLL fusion proteins aberrantly activate the expression of HSC-specific genes remain elusive.

MLL fusion proteins form a complex with MENIN (Yokoyama *et al*, 2005; Yokoyama *et al*, 2004), which leads to further association with LEGDF (Yokoyama & Cleary, 2008). MLL fusion proteins bind to their target chromatin through the CXXC domain of MLL, which specifically recognizes unmethylated CpGs, and the PWWP domain of LEDGF, which selectively binds to di/tri-methylated histone H3 lysine 36 (H3K36me2/3) (Okuda *et al*, 2014). The CXXC and PWWP domains constitute the minimum targeting module (MTM) which can stably bind to the MLL target gene promoters (e.g., *HOXA9*). Because unmethylated CpGs and H3K36me2/3 marks are associated with transcriptional activation, MLL fusion proteins target a broad range of previously transcribed CpG-rich promoters. Although MLL fuses with a variety of partners, most *MLL*-rearranged leukemia cases are caused by fusions with the AF4 family (e.g., AF4 also known as AFF1, AF5Q31 also known as AFF4) and ENL family (e.g., ENL also known as MLLT1, AF9 also known as MLLT3) (Meyer *et al*., 2018). AF4 family proteins form a biochemically stable complex with ENL family proteins and the P-TEFb elongation factor (composed of CDK9 and CyclinT1/2), which we termed as AEP (as in AE4 family/ ENL family/ P-TEFb complex) (Yokoyama *et al*, 2010). AF4 family proteins recruit the SL1 complex (Okuda *et al*, 2015), which is composed of TBP and TAF1A/B/C/D subunits, and is known to initiate ribosomal RNA transcription by RNA polymerase I (Goodfellow & Zomerdijk, 2013). An artificial construct of MTM fused to the binding platform for SL1 activated *Hoxa9* and transformed hematopoietic progenitors (HPCs) (Okuda *et al*., 2015), indicating that AF4 activates RNA polymerase II (RNAP2)-dependent transcription, presumably by loading TBP onto the target promoters via SL1. However, why MLL fusion proteins preferentially use the AEP/SL1-mediated transcription pathway is unclear.

In this study, we identified the evolutionarily conserved TRX2 domain of MLL as a key structure required for aberrant self-renewal mediated by MLL fusion proteins. Subsequent proteomic approach identified the HBO1 (also known as KAT7 and MYST2) HAT complex as an associating factor for the TRX2 domain. HBO1 is a member of MYST HAT family responsible of the bulk of Histone H3 lysine 14 acetylation (H3K14ac) (Mishima *et al*, 2011), and was recently identified as a therapeutic vulnerability of leukemia stem cells by genetic screening (Au *et al*, 2020; MacPherson *et al*, 2020). However, its molecular functions on leukemic proteins remain largely elusive. Our detailed structure/function analysis demonstrated that HBO1-MLL interaction promotes AEP-dependent gene activation in MLL fusion-mediated leukemic transformation. Moreover, another leukemic fusion of nucleoporin-98 (NUP98) and HBO1(i.e. NUP98-HBO1) also transformed HPCs via association with MLL. Hence, we propose that HBO1-MLL interaction modules can be utilized as molecular targets for developing drugs that specifically dismantle the oncogenic transcriptional machinery.

## Results

### TRX2 domain-mediated functions promote MLL fusion-dependent leukemic transformation

MLL fusion proteins confer unlimited self-renewal ability, leading to immortalization of HPCs (Lavau *et al*, 1997). *MLL-ELL* is one of the frequently observed MLL fusions associated with acute myelogenous leukemia (AML) (Meyer *et al*., 2018). To determine the domain structures required for MLL-ELL-mediated leukemic transformation, we performed myeloid progenitor transformation assays (Lavau *et al*., 1997; Okuda & Yokoyama, 2017b), wherein murine HPCs were retrovirally transduced with an MLL fusion gene, and cultured in semisolid medium supplemented with myeloid cytokines (Figure 1A). MLL-ELL transformed HPCs, as previously reported (DiMartino *et al*, 2000; Luo *et al*, 2001), featuring vigorous colony forming capacities at the third and fourth passages, and elevated *Hoxa9* expression at the first and second passages, whereas a deletion mutant lacking the TRX2 domain failed to transform (Figures 1A and S1A), underscoring the biological significance of the TRX2 domain. A minimalistic artificial construct of HA-tagged MTM fused to an intact Occludin-homology domain (OHD) of ELL failed to transform HPCs (see MTMh-ELĹ’), whereas inclusion of the TRX2 domain to MTM (hereafter denoted as MTMT) caused constitutive activation of *Hoxa9* and immortalization of HPCs (see MTMTh-ELĹ’). These results indicate that MLL-ELL transforms HPCs through TRX2 domain-mediated functions.

**Figure 1.**
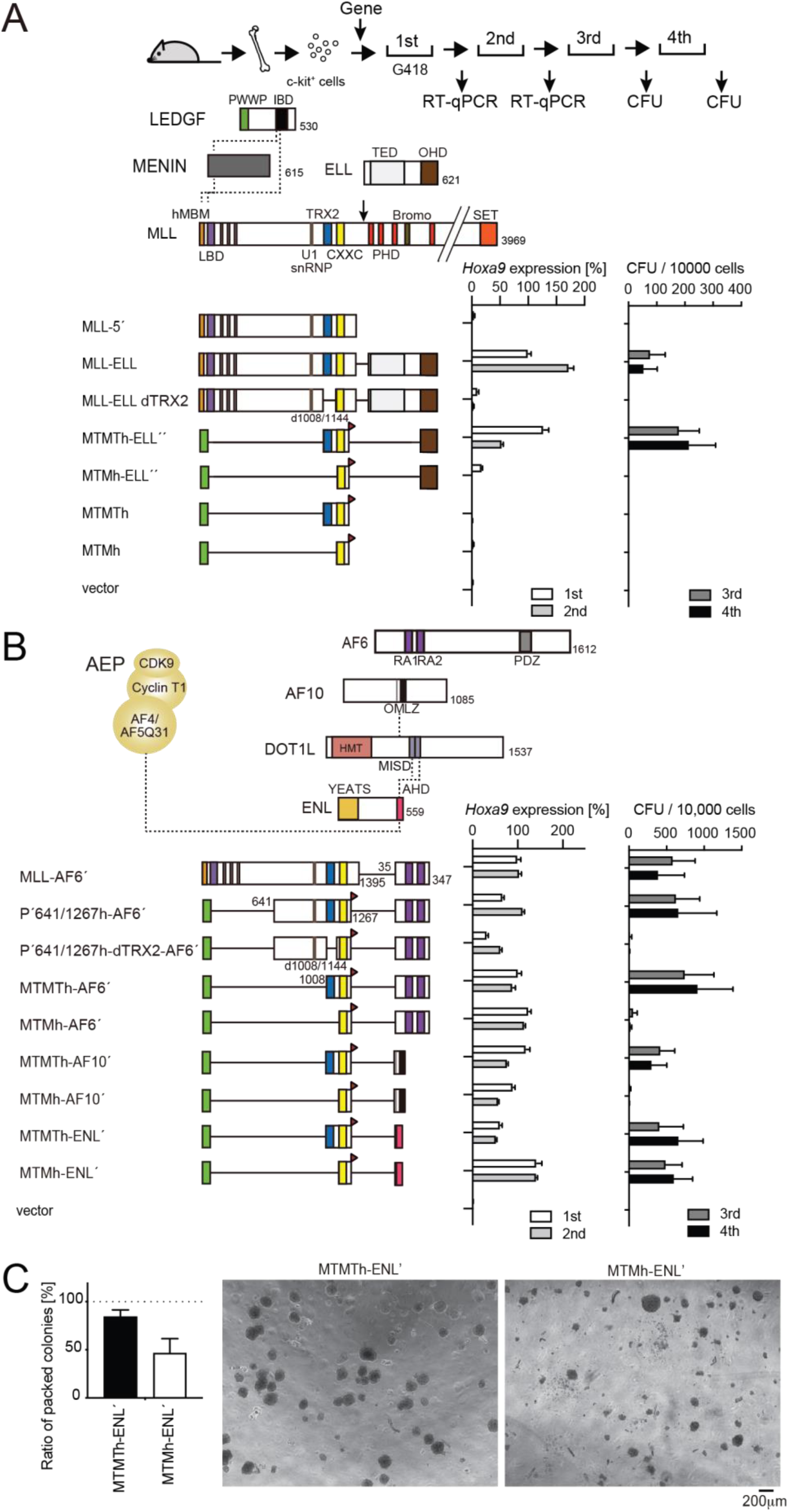
TRX2 domain-mediated functions promote MLL-fusion-dependent leukemic transformation. A. Structure/function analysis of MLL-ELL. Various MLL-ELL constructs were examined for transformation of myeloid progenitors. HA-tag (h: indicated as a red triangle) was fused to MTM and MTMT constructs. A schema of myeloid progenitor transformation assay is shown on top. Hoxa9 expression normalized to Gapdh in first round and second round colonies (left) is shown as the relative value of MLL-ELL (arbitrarily set at 100%) with error bars (mean ± SD of PCR triplicates). Colony-forming ability at third- and fourth-round passages (right) is shown with error bars (mean ± SD of ≥3 biological replicates). B. Requirement of the TRX2 domain for various MLL fusion proteins in leukemic transformation. Various MLL fusion constructs were examined for transformation of myeloid progenitors as in Figure 1A. C. Colony morphologies of MTMTh or MTMh-ENĹ-transformed cells. The colonies on day 5 of fourth passage are shown with a scale bar. The ratio of compact colonies (≥100 total colonies were counted in each experiment) is shown on the left (mean ± SD of six biological replicates). Representative images are shown on the right.

Next, we tested the structural requirements of other MLL fusions (i.e., MLL-AF6, MLL-AF10, and MLL-ENL) (Figures 1B and S1A). An MLL-AF6 fusion construct, in which MLL is fused to the RA1 and RA2 domains of AF6 (see MLL-AF6’), fully transformed HPCs as previously reported (Liedtke *et al*, 2010). The minimalistic MTM-AF6 fusion construct (see MTMh-AF6’) activated *Hoxa9* expression in the early passages but produced modest numbers of colonies in later passages, while inclusion of the TRX2 domain (see MTMTh-AF6’) conferred much more vigorous transforming capacities. Deletion of the TRX2 domain from a PWWP-MLL-AF6 fusion construct containing the residues 641/1267 of MLL (P’641/1267h-AF6’) abrogated its transforming ability, suggesting that MLL-AF6 requires the TRX2 domain to exert its full transforming potential (Figures 1B and S1A). Nonetheless, an MLL-AF6’ construct lacking the TRX2 domain immortalized HPCs albeit less efficiently compared to MLL-AF6’ (Figure S1B, see MLL-AF6’ dTRX2), suggestive of some compensatory functions mediated by the MLL structure retained in MLL-AF6’ but missing in P’641/1267h-AF6’ (i.e., the residues 1/640 and 1268/1395). MLL-AF10 showed a similar trend, in line with our previous report (Okuda *et al*, 2017) (Figures 1B and S1A, B). On the other hand, the oncogenic properties of MLL-ENL were not severely affected by the loss of TRX2 domain in terms of colony forming capacity (Okuda *et al*., 2014). However, the colony morphology of immortalized cells (ICs) transformed by the MTMh-ENL construct (MTMh-ENĹ-ICs) was more differentiated compared to that of MTMTh-ENĹ-ICs (Figure 1C), suggesting that the TRX2 domain is required to block differentiation. Leukemogenesis in vivo was compromised by deletion of the TRX2 domain for MLL-AF6, -AF10, and -ENL (Figure S1C). These results indicate that MLL fusion proteins rely on TRX2 domain-mediated functions for leukemic transformation to varying degrees depending on their fusion partners. Among the MLL fusions tested, MLL-ELL most heavily relies on TRX2 domain-mediated functions.

### HBO1 complex associates with MLL proteins via the TRX2 domain at promoters

We previously showed that the TRX2 domain binds to AF4 family proteins (Okuda *et al*., 2017). Indeed, a FLAG-tagged MLL construct encompassing the residues 869/1152 efficiently co-precipitated with exogenously expressed AF4 or AF5Q31 (Figure S2A, see fMLL 869/1152+37aa). However, deletion of the TRX2 domain from the FLAG-tagged MLL-5’construct (containing the residues 1/1395: fMLL-5’) did not impair co-precipitation of AF4, indicating that the interaction with AF4 is not mediated by the TRX2 domain. Sequencing analysis of the vector constructs revealed that the fMLL 869/1152+37aa construct contained an additional coding sequence for 37 residues derived from the pCMV5 vector tethered in frame, which corresponds to a part of the Chorionic somatomammotropin hormone 1 gene (Figure S2B). Removal of the additional 37 residues by introducing a stop codon resulted in complete loss of association with AF4 family proteins (Figure S2A, see fMLL 869/1251). Moreover, a FLAG-tagged GAL4 fusion construct tethered to the additional 37 residues co-precipitated with AF5Q31 (see fGAL4+37aa). Thus, we concluded that our previous claim for the TRX2 domain as a binding platform for AF4 family proteins was false.

To identify bona fide associating factors for the TRX2 domain, we purified two exogenously expressed TRX2 domain-containing proteins (i.e., fGAL4-MLL 869/1124 and MTMTh) from the chromatin fraction of HEK293T cells using the fractionation-assisted chromatin immunoprecipitation (fanChIP) method (Okuda *et al*., 2015), and analyzed by mass spectrometry (Figure 2A). Components of the HBO1 complex including HBO1, PHF16 (also known as JADE3), MEAF6, and ING5 were detected in the purified materials (Avvakumov *et al*, 2012). Immunoprecipitation (IP)-western blotting analysis confirmed that fGAL4-MLL 869/1124 co-precipitated with HBO1 complex components but not with exogenously expressed AF4 (Figures 2B and S3A). A series of MTMT fusion proteins associated with HBO1 and PHF16, while the respective MTM fusion proteins did not (Figure 2C), confirming that the TRX2 domain mediates interaction with the HBO1 complex. Domain mapping analysis of MLL showed that the residues 869/1124 contain the major binding domains for the HBO1 complex (Figures S3A). Moreover, fGAL4-MLL 1052/1124 co-precipitated with the HBO1 complex components after DNaseI treatment, indicating that association of MLL and HBO1 is not mediated by DNA (Figure S3B). It should be noted that a small amount of HBO1 co-precipitated with the MLL proteins lacking the TRX2 domain (Figure S3C, see fMLL-5’ dTRX2 and fMLL-ENL dTRX2), suggesting that there is a secondary binding domain for the HBO1 complex outside of the TRX2 domain, which may account for the moderate effects of TRX2 domain deletion of the full length MLL fusion constructs (Figure S1B).

**Figure 2.**
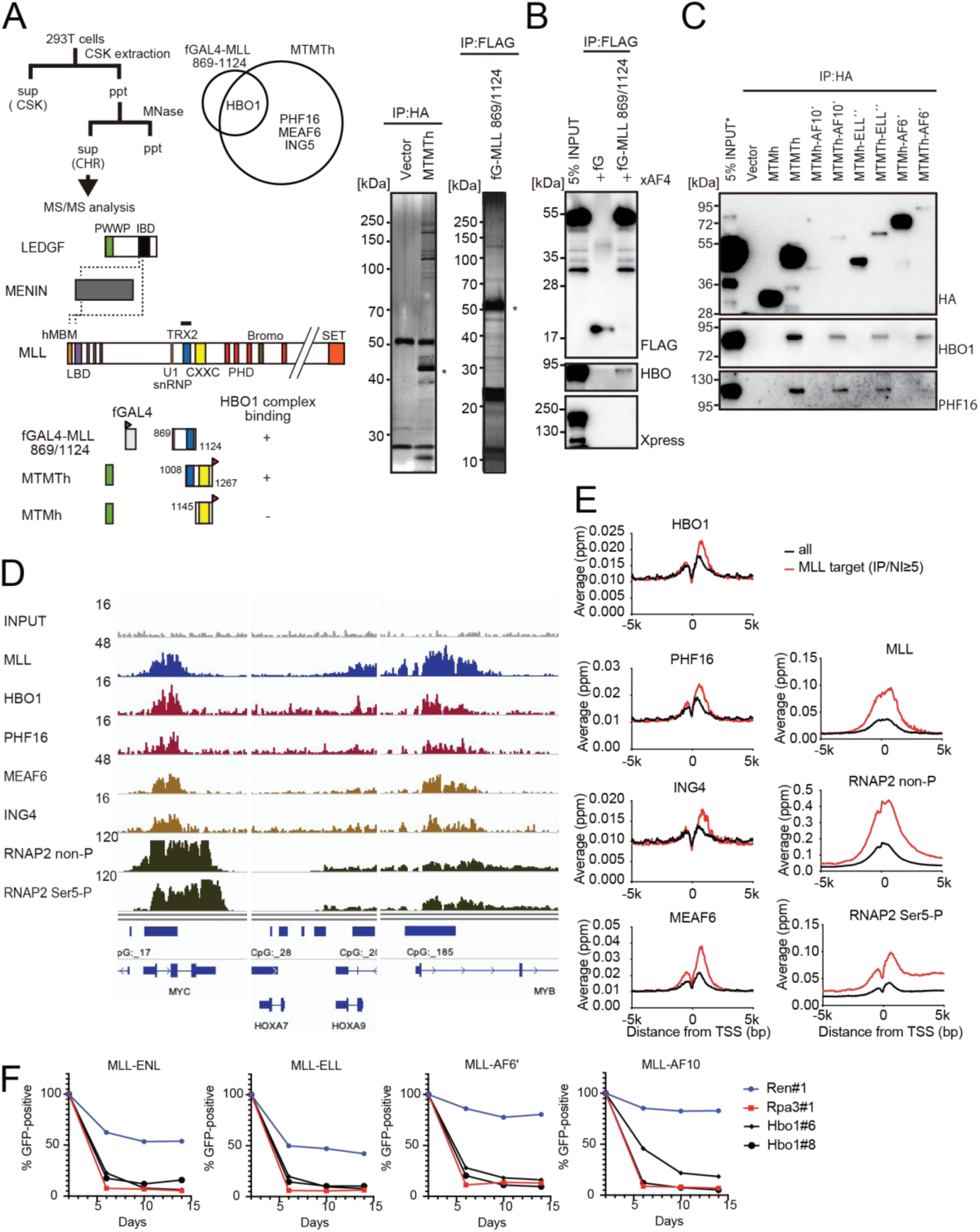
HBO1 complex associates with MLL proteins via the TRX2 domain at promoters. A. Purification of TRX2 domain-associating factors. A FLAG-tagged (f: indicated as a blue triangle) GAL4 DNA binding domain fused with the MLL fragment containing the residues 869-1124 or HA-tagged MTMT fragment was transiently expressed in HEK293T cells. A schema of fanChIP method is shown on top. The transgene products were purified from the chromatin fraction and analyzed by mass spectrometry. Silver-stained images (right) of the purified materials are shown. Asterisk indicates the position of the transgene products. A Venn diagram of identified TRX2 domain-associating factors by mass spectrometry is shown. B. Association of GAL4-TRX2 fusion with HBO1. IP-western blotting of the chromatin fraction of HEK293T cells transiently expressing FLAG-tagged GAL4-MLL 869/1124 construct and Xpress-tagged AF4 (xAF4) was performed. Co-purification of HBO1, but not xAF4, was confirmed. C. TRX2 domain-dependent association with the HBO1 complex. IP-western blotting of the chromatin fraction of virus-packaging cells, transiently expressing various HA-tagged MTMT (or MTM) fusion constructs, was performed. D. Genomic localization of MLL and the HBO1 complex components in HB1119 cells. The ChIP-seq profiles were visualized using the Integrative Genomics Viewer (The Broad Institute). The minimum value of the y-axis was set at 0, while the maximum value for each sample is indicated. E. Average distribution of proteins near the TSSs of HB1119 cells. Genes whose MLL ChIP signal/input ratio at the promoter proximal transcribed region was ≥ 5 were defined as MLL target genes. Average ChIP signal distribution of indicated proteins at the MLL target genes (red) or all genes (black) is shown. The y-axis indicates the frequency of the ChIP-seq tag count (ppm) in 25 bp increments. F. Requirement of HBO1 for myeloid progenitors immortalized by various oncogenes. sgRNA competition assays for *Hbo1* were performed on immortalized myeloid progenitors. The ratio of GFP-positive cells co-expressing sgRNA was measured by flow cytometry. sgRNA for Renilla luciferase (Ren) was used as a negative control, which does not affect proliferation. sgRNA for Rpa3 was used as a positive control, which impairs proliferation.

Interaction between endogenous MLL proteins and HBO1 was confirmed in leukemia cell lines such as HB1119 and REH (Figure S3D, E). Chromatin immunoprecipitation (ChIP) followed by deep sequencing (ChIP-seq) of HB1119 cells, which endogenously express MLL-ENL, demonstrated that HBO1 complex components colocalized with MLL-ENL at the MLL target genes (e.g. *MYC*, *HOXA9* and *MYB*) in a genome-wide manner (Figure 2D, E). CRISPR/Cas9-mediated sgRNA competition assays demonstrated that *Hbo1* is required for the proliferation of various MLL fusion-ICs (Figure 2F). These results are consistent with the recent reports, which showed that the HBO1 complex associates with multiple MLL fusion proteins, and plays a critical role in the maintenance of leukemia stem cells (Au *et al*., 2020; MacPherson *et al*., 2020). Taken together, MLL fusion proteins associate with the HBO1 complex through the TRX2 domain at the target promoters.

### MLL recruits the HBO1, AEP, and SL1 complexes to promoters

To elucidate the functional relationship between wild type MLL and the HBO1 complex, we examined the genomic localization of the HBO1 complex, wild type MLL, and its related transcriptional regulators in HEK293T cells (Figure 3A). MLL has been shown to localize at the transcription start sites (TSSs) of CpG-rich genes including, *RPL13A*, *MYC* and *CDKN2C* (Okuda *et al*., 2017). Distribution of the HBO1 complex was enriched at promoter-proximal transcribed regions (0-2 kb from the TSS) (Figure 3B) consistent with a previous report (Avvakumov *et al*., 2012), suggesting its implication in transcription initiation/elongation. Similarly as in HB1119 cells, the HBO1 complex was localized at the MLL-occupied promoters in a genome-wide manner. We previously showed that MLL associates with the MOZ complex and yet-to-be activated RNAP2 whose heptapeptide repeats are not phosphorylated (RNAP2 non-P) on target promoters (Miyamoto *et al*, 2020), and colocalizes with various ENL-containing complexes including the AEP and DOT1L complexes (Okuda *et al*., 2017). The ChIP signal intensities of MLL were highly correlated with those of HBO1, MOZ, and RNAP2 non-P (Figure 3C), supporting the functional interactions of MLL with those MLL-associated factors. The ChIP signals of AEP (e.g., AF4, ENL) and SL1 (e.g., TAF1C) are weakly corelated with that of MLL, presumably because AEP and SL1 are indirectly recruited to MLL target promoters in a context dependent manner.

**Figure 3.**
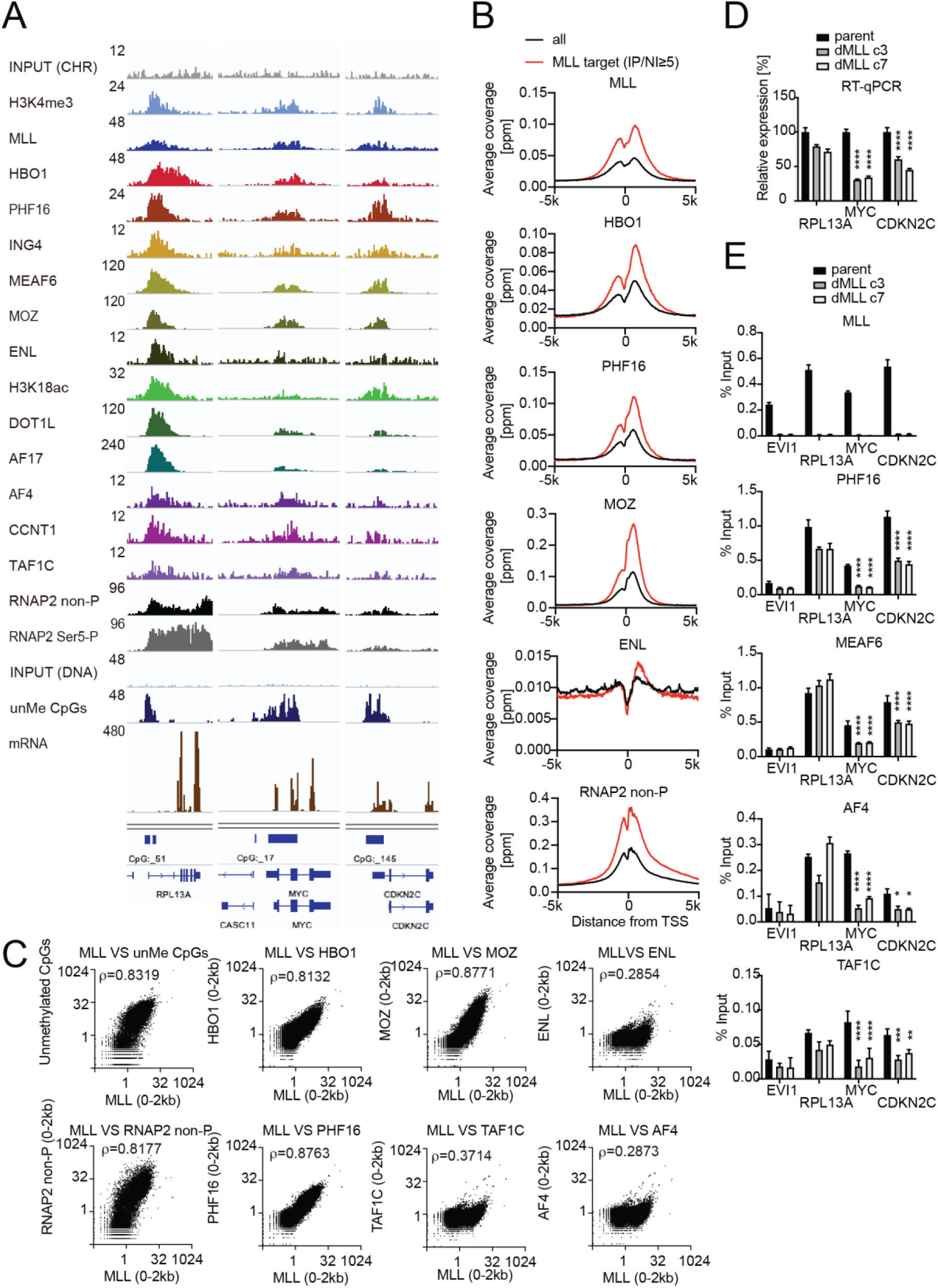
MLL recruits the HBO1 and AEP/SL1 complexes to promoters. A. Genomic localization of various transcriptional regulators/epigenetic marks in HEK293T cells. ChIP-seq analysis was performed on the chromatin of HEK293T cells for the indicated proteins/modifications. CIRA-seq data for unmethylated CpGs (unMe CpGs) and RNA-seq data are shown for comparison. B. Average distribution of proteins near TSSs of HEK293T cells. Average ChIP signal distribution of indicated proteins at the MLL target genes (red) or all genes (black) is shown as in Figure 2E. C. Relative occupation by multiple factors at all TSSs. ChIP-seq or CIRA-seq tags of indicated proteins at all genes were clustered into a 2 kb bin (0 to +2 kb from the TSS) and are presented as XY scatter plots. Spearman’s rank correlation coefficient (ρ) is shown. D. Expression of MLL target genes in two independently generated *MLL*-deficient clones. Relative expression levels normalized to *GAPDH* are shown with error bars (mean ± SD of PCR triplicates) by RT-qPCR. Data are redundant with our previous report(Miyamoto *et al*., 2020) E. Localization of MLL, the HBO1 complex, AEP, and the SL1 complex in two independently generated *MLL*-deficient clones. ChIP-qPCR was performed for indicated genes using qPCR probes designed for the TSS of each gene. ChIP signals are expressed as the percent input with error bars (mean ± SD of PCR triplicates). Statistical analysis was performed by ordinary two-way ANOVA comparing each sample with the parent cells. \**P* ≤ 0.05., \*\**P* ≤ 0.01. \*\*\**P* ≤ 0.001., \*\*\*\**P* ≤ 0.0001.

To examine the role of MLL in the genomic localization of the HBO1 complex, we analyzed *MLL*-deficient HEK293T cells by qRT-PCR and ChIP-qPCR. *MLL* knockout reduced the expression of *MYC* and *CDKN2C* (Figure 3D), as previously reported (Miyamoto *et al*., 2020), and caused a marked reduction in the ChIP signals of HBO1 complex components (i.e., PHF16, MEAF6) at the *MYC* and *CDKN2C* loci (Figure 3E). In addition, the ChIP signals of AF4 and TAF1C were also reduced at these loci. These results indicate that MLL recruits the HBO1, AEP, and SL1 complexes to the target promoters.

### MLL-ELL transforms through the common binding platform for AF4 and EAF1

Next, we investigated the function of the ELL portion in MLL-ELL-mediated leukemic transformation. ELL has a transcriptional elongation activity, and is associated with a variety of proteins including AF4- and EAF-family proteins (Lin *et al*, 2010; Shilatifard *et al*, 1996; Simone *et al*, 2003; Simone *et al*, 2001). Myeloid progenitor transformation assays demonstrated that an intact OHD, which is responsible for association with both AF4 and EAF1 (Lin *et al*., 2010; Simone *et al*., 2001), is required for transformation as previously reported (Figures 4A, B and S4A) (DiMartino *et al*., 2000; Luo *et al*., 2001). ChIP-qPCR analysis of FLAG-tagged GAL4 constructs fused to ELL confirmed that OHD is responsible for the recruitment of both AF4 and EAF1 (Figure 4C). Deletion of OHD resulted in loss of interaction with both AF4 and EAF1 family proteins, which is correlated with the transforming properties (Figure 4A-D). It should be noted that several processed forms of AF4 (e.g., 110kDa) were observed in the co-precipitates, whose amounts were more abundant in the chromatin fraction of HEK293T cells transiently expressing GAL4-ELL proteins harboring an intact OHD (Figure 4B, D), suggesting that ELL tethers both processed and unprocessed forms of AF4 to the chromatin. ELL-AF4 interaction was attenuated by co-expression of EAF1, whereas ELL-EAF1 interaction was augmented by co-expression of AF4 (Figures 4E, F), suggesting that these interactions occur sequentially, wherein the ELL-AF4 interaction precedes the ELL-EAF1 interaction (Figure 4G). Taken together, MLL-ELL exerts its transforming properties through the common binding platform for AF4 and EAF1.

**Figure 4.**
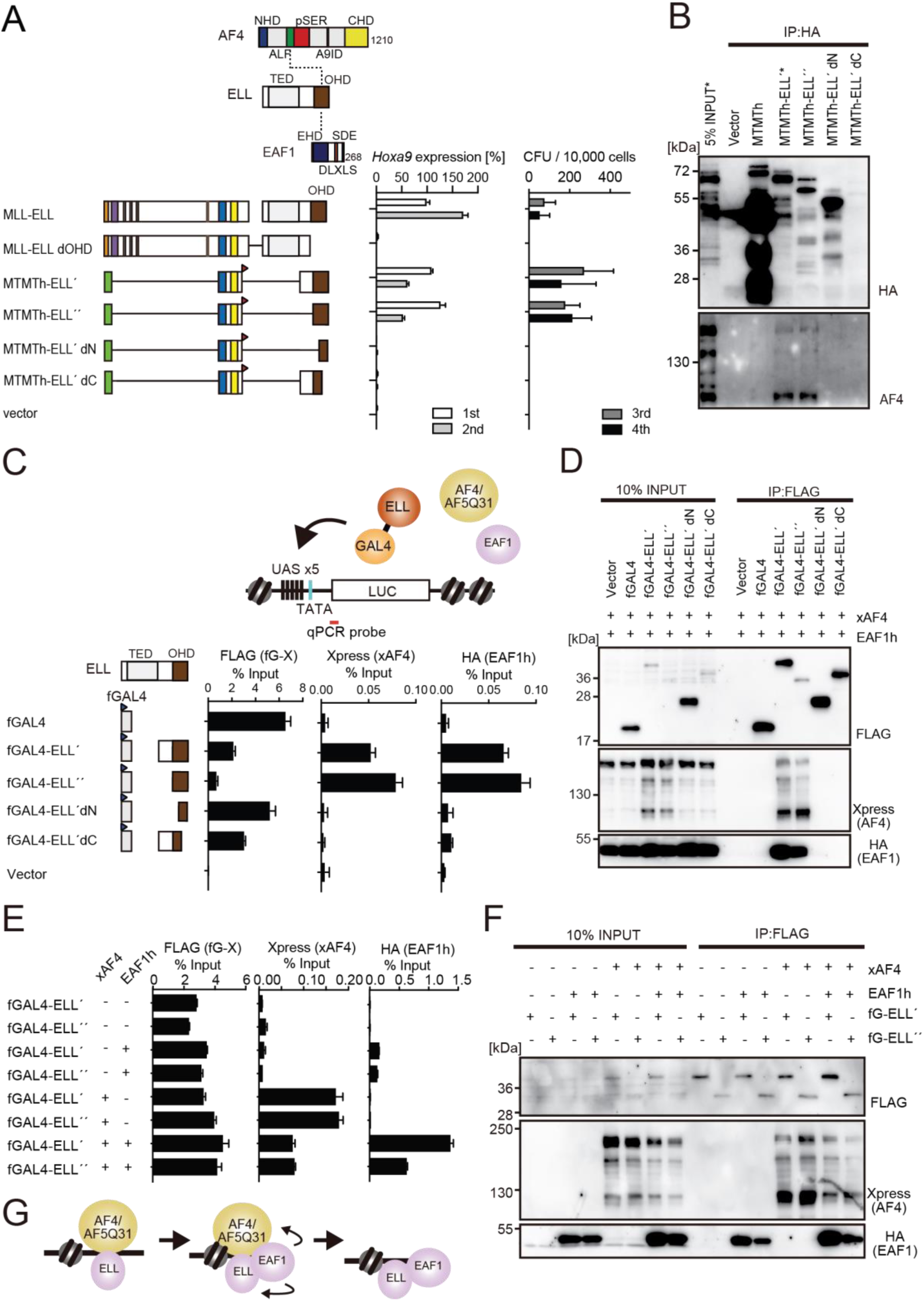
MLL-ELL transforms through the common binding platform for AF4 and EAF1. A. Requirement of the OHD in MLL-ELL-mediated transformation. Various MLL-ELL constructs carrying mutations in the ELL portion were examined for transformation of myeloid progenitors, as in Figure 1A. B. OHD-dependent association with AF4. IP-western blotting of the chromatin fraction of virus packaging cells, transiently expressing various HA-tagged MTMT-ELL fusion constructs, was performed. Endogenous AF4 proteins co-purified with MTMTh-ELL proteins were visualized by anti-AF4 antibody. C. Recruitment of exogenously expressed AF4 or EAF1 by ELL. HEK293TL cells (Okuda *et al*., 2015), which harbor GAL4-responsive reporter, were transfected with FLAG-tagged GAL4 fusion proteins, xAF4, and HA-tagged EAF1 (EAF1h), and were subjected to ChIP-qPCR analysis. A qPCR probe near the GAL4-responsive elements (UAS) was used. The ChIP signals were expressed as the percent input with error bars (mean ± SD of PCR triplicates). TATA: TATA box, LUC: Luciferase. D. Association of ELL with exogenously expressed AF4 or EAF1 on chromatin. IP-western blotting of the chromatin fraction of HEK293TL cells transiently expressing various FLAG-tagged GAL4-ELL proteins along with xAF4 and EAF1h was performed. E. Effects of overexpression of AF4 or EAF1 on ELL complex formation at the promoter. ChIP-qPCR analysis of HEK293TL cells transiently expressing various combinations of FLAG-tagged GAL4-ELL proteins, xAF4, and EAF1h was performed as in Figure 4C. F. Effects of overexpression of AF4 or EAF1 on ELL complex formation. IP-western blotting of the chromatin fraction of HEK293TL cells, transiently expressing various combinations of FLAG-tagged GAL4-ELL proteins, xAF4, and EAF1h, was performed as in Figure 4D. G. A model of sequential association between ELL, AF4 family proteins, and EAF family proteins.

### AF4 and EAF1 form two distinct SL1/MED26-containing complexes

AF4 and EAF1 family proteins share some structural similarities; both have the SDE motif enriched with serine, aspartic acid, and glutamic acid, and the DLXLS motif whose consensus sequence is LXXDLXLS (Figure 5A, B). The NKW motif was found in the AF4 family but not in the EAF family. It has been demonstrated by our group and others that the SDE motif of AF4 associates with the SL1 complex (Okuda *et al*., 2015) and its DLXLS motif associates with MED26 (Okuda *et al*, 2016; Takahashi *et al*, 2011). IP-western blotting analysis demonstrated that EAF1 also associated with SL1 and MED26 through its C-terminal domain containing the SDE and DLXLS motifs (Figure 5C). ChIP-qPCR analysis confirmed that the C-terminal portion of EAF1 recruited TAF1C and MED26 to the GAL4-responsive promoter (Figure 5D, see fGAL4-EAF1-C). Nevertheless, the GAL4-ELL fusion failed to associate with or recruit TAF1C and MED26 (Figure 5C, D, see fGAL4-ELĹ), indicating that ELL-bound EAF1 or AF4 is unable to interact with TAF1C and MED26, and therefore must dissociate from ELL to form a complex with SL1 and MED26. It should be noted that GAL4-EAF1 failed to pull down endogenous AF4 and ENL, while GAL4-AF4 containing an ELL binding domain (ALF) (see fGAL4-AF4-2N&C) also failed to pull down exogenously expressed EAF1, indicating that the AF4/ELL/EAF1 trimer complex is unstable and that ELL mostly binds AF4- or EAF-family proteins in a mutually exclusive manner.

**Figure 5.**
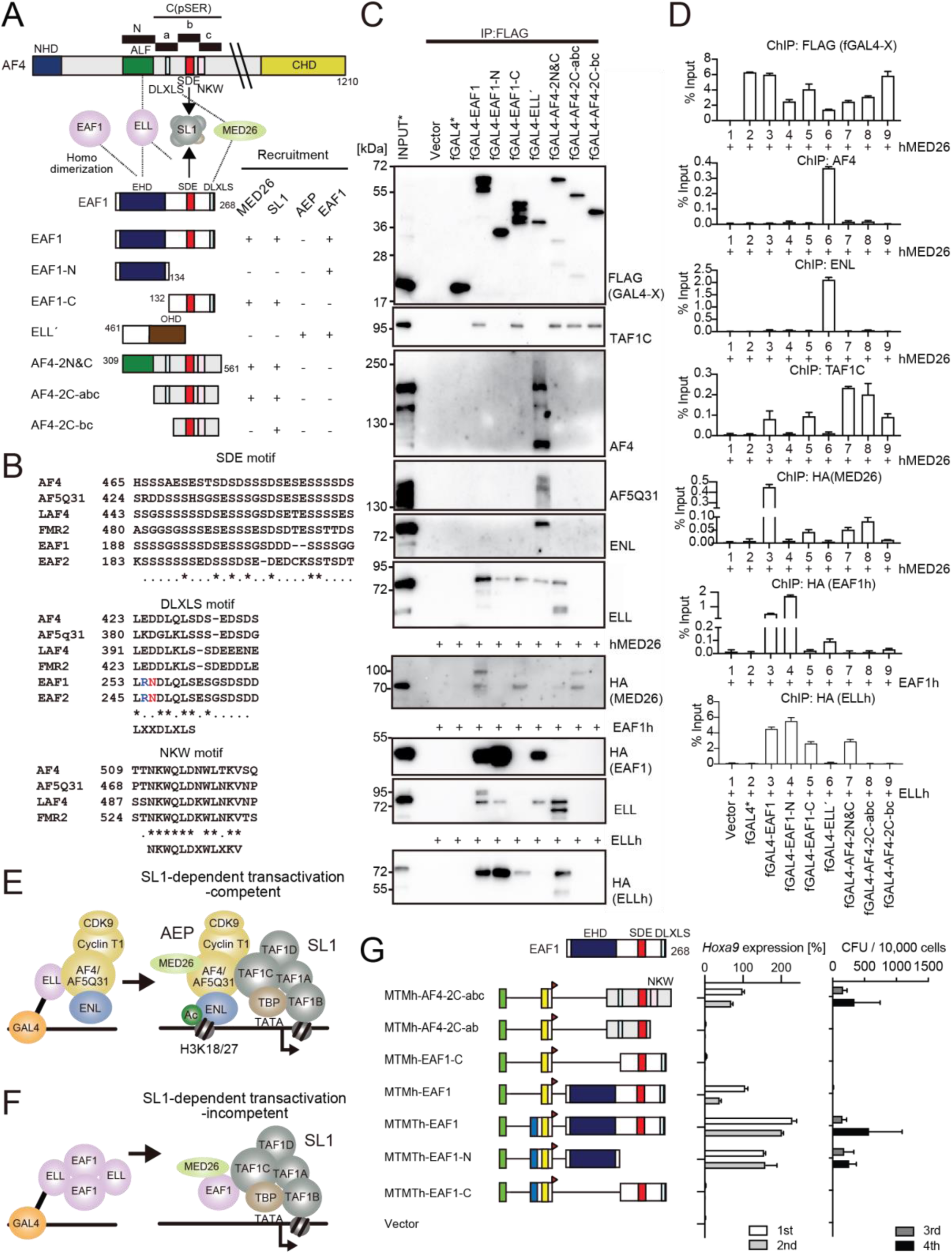
AF4 and EAF1 form distinct SL1/MED26-containing complexes. A. A schema of the structures of AF4, ELL, and EAF1. NHD: N-terminal homology domain, ALF: AF4/LAF4/FMR2 homology domain responsible for interaction with ELL, pSER: poly serine, DLXLF: DLXLF motif, SDE: SDE motif responsible for interaction with SL1, DLXLS: DLXLS motif responsible for interaction with MED26, CHD: C-terminal homology domain, EHD: EAF homology domain responsible for homodimerization and interaction with ELL. B. Alignment of the amino acid sequences of the SDE, DLXLS, and NKW motifs of human AF4 family proteins and EAF family proteins. Conserved residues are indicated by asterisks. C. Association of EAF1, ELL, and AF4 domains with various associating factors on chromatin. IP-western blotting of the chromatin fraction of HEK293TL cells transiently expressing various FLAG-tagged GAL4 fusion proteins, with or without indicated HA-tagged constructs, was performed as in Figure 4D. Endogenous proteins were detected by specific antibodies for each protein, while exogenous proteins were detected by antibodies for FLAG or HA tag. D. Recruitment of various transcriptional regulatory proteins by EAF1, ELL, and AF4 domains. ChIP-qPCR analysis of HEK293TL cells transiently expressing various combinations of FLAG-tagged GAL4 fusion proteins along with indicated HA-tagged constructs was performed as in Figure 4C. E. Putative complex recruitment mediated by ELL-AF4 interaction. F. Putative complex recruitment mediated by ELL-EAF1 interaction. G. Structural requirement of MTM-AF4 pSER fusion and MTMT-EAF1 fusion. Various MTMh/MTMTh constructs fused with EAF1 domains were examined for transformation of myeloid progenitors as in Figure 1A.

In addition, GAL4-EAF1 co-precipitated with exogenously expressed EAF1 through the EAF family homology domain (EHD), indicating that EAF1 forms a homodimer (Figure 5C, D). It has been shown that EAF1 binds ELL through two distinct contacts (Simone *et al*., 2003). Accordingly, both N- and C-terminal halves of EAF1 co-precipitated with endogenous ELL (see fGAL4-EAF1-N and -C, where no HA-tagged protein was co-expressed). However, when EAF1 was overexpressed, co-precipitation of endogenous ELL by the C-terminal half of EAF1 was no longer detected, suggesting that free ELL preferentially binds to an EAF1 dimer over the single contact-mediated interaction with the C-terminal half of EAF1. GAL4-ELL co-precipitated with endogenous ELL presumably mediated by an EAF1 dimer, whereas it failed to pull down ELL when exogenous ELL was overexpressed to absorb free EAF1 dimers. Taken together, these results suggest that ELL forms at least two different stable complexes, one is with AF4 family proteins, which subsequently leads to the formation of an AEP/SL1/MED26 complex (Figure 5E), and the other is an EAF1/ELL dimer, which leads to the formation of an EAF1/SL1/MED26 complex (Figure 5F). These two SL1/MED26 containing complexes are similar in composition, but different in three key functions. The AEP/SL1/MED26 complex has the NKW motif which is essential for AF4-dependent gene activation (Okuda *et al*., 2015), and contains P-TEFb which promotes transcription elongation, and ENL family proteins which tether AEP on acetylated chromatin (Erb *et al*, 2017; Li *et al*, 2014; Wan *et al*, 2017). Thus, we presumed that the AEP/SL1/MED26 complex is competent for transactivation whereas the EAF1/SL1/MED26 complex is not.

### Interaction with AEP drives MLL-ELL-mediated leukemic transformation, while homodimerization promotes MLL-EAF1-mediated transformation

Previously, Luo et al. demonstrated that an artificial fusion construct of MLL and EAF1 transformed HPCs and induced leukemia (Luo *et al*., 2001), which suggested that the ELL-EAF1 interaction played a major role in MLL-ELL-mediated leukemic transformation. To examine the structural requirements for MLL-EAF1-mediated transformation, we generated artificial fusion constructs in which MTM or MTMT is fused to EAF1 domains and examined their transforming properties (Figures 5G and S5B). As reported previously (Okuda *et al*., 2015), an MTM construct fused to the AF4 portion containing the SDE and NKW motifs (see MTMh-AF4-2C-abc) activated *Hoxa9* expression and immortalized HPCs, while removal of the NKW motif (see MTMh-AF4-2C-ab) resulted in loss of transformation (Figure 5G). Accordingly, an MTM construct fused to the C-terminal half of EAF1 containing the SDE motif but lacking the NKW motif failed to transform HPCs (see MTMh-EAF1-C). However, an MTM construct fused to the entire EAF1 demonstrated partial transforming properties, which maintained the expression of *Hoxa9* in the early passages, but failed to immortalize HPCs (see MTMh-EAF1). MTMT constructs fused to the EHD fully transformed HPCs (Figure 5G, see MTMTh-EAF1 and MTMTh-EAF1-N). This trend is reminiscent of MLL-AF6 which transforms HPCs by homodimerization (Figure 1B) (Liedtke *et al*., 2010), and suggests that MLL-EAF1 transforms by EHD-mediated homodimerization, rather than by ELL-EAF1 interaction. Knockdown of *Enl* in MLL-ELL-ICs perturbed colony formation and *Hoxa9* expression, while its effects on MLL-ENL-ICs were relatively limited presumably because MLL-ENL can directly recruit AEP. With these results, we speculated that MLL-ELL transforms HPCs through recruitment of AEP, while MLL-EAF1 transforms via homodimerization in a mechanism similar to MLL-AF6.

### MLL-ELL transforms hematopoietic progenitors via association with AEP, but not with EAF1 or p53

To further evaluate the roles of the ELL-AF4 and ELL-EAF1 interactions in MLL-ELL-mediated leukemic transformation, we next introduced S600A/K606T double mutation (hereafter denoted as SA/KT) to the ELL portion, which was initially predicted to impair ELL-AF4 interaction based on the structural data for the ELL2/AF5Q31 complex (Qi *et al*, 2017). Indeed, the mutation severely attenuated ELL-AF4 interaction (Figure 6A, see fGAL4-ELĹ’ SA/KT). However, a substantial amount of AF4 (mostly the processed from of AF4) remained associated to the SA/KT mutant, while co-precipitation of EAF1 was completely abolished. These results suggest that SA/KT mutation abolishes the primary contact of ELL for AF4 and EAF1, while there is a secondary contact for AF4 which is unaffected by this mutation. Western bloting of the input samples of the chromatin fraction showed that fGAL4-ELĹ’ SA/KT increased the amount of processed forms of AF4 in the chromatin fraction like fGAL4-ELĹ’, thus confirming the interaction with fGAL4-ELĹ’ SA/KT and AF4. ChIP-qPCR analysis confirmed that exogenously expressed AF4 was recruited to the target chromatin by fGAL4-ELĹ’ SA/KT, while EAF1 was not (Figure 6B). Moreover, fGAL4-ELĹ’ SA/KT co-precipitated endogenous AF4 and AF5Q31, while it failed to pull down p53, another ELL associating factor (Wiederschain *et al*, 2003) (Figure S6A). fGAL4-ELĹ’ SA/KT recruited a substantial amount of endogenous ENL and CyclinT1 to the GAL4-responsive promoter (Figure S6B), indicating that the SA/KT mutant is competent for loading AEP onto chromatin.

**Figure 6.**
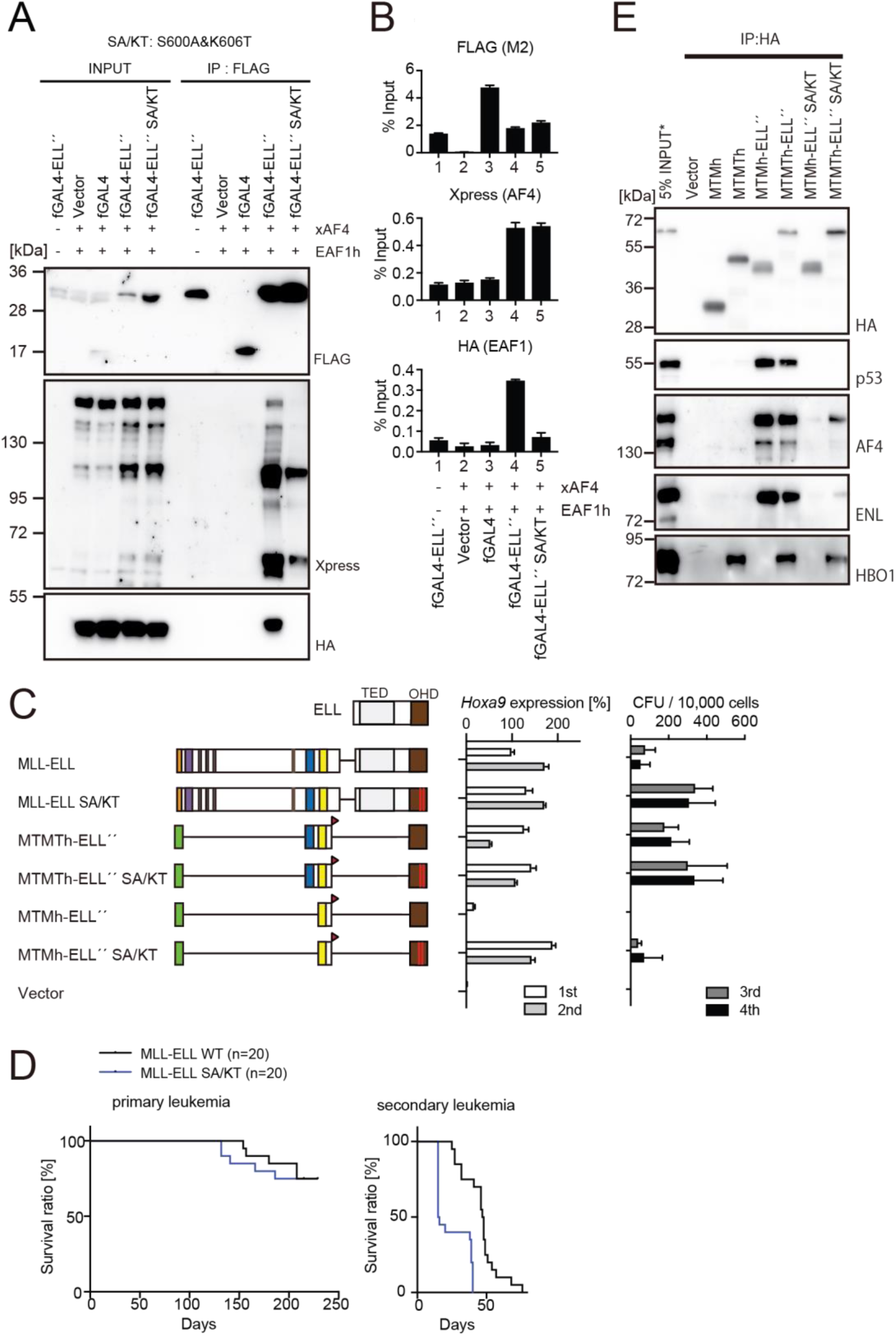
MLL-ELL transforms hematopoietic progenitors via association with AEP, but not with EAF1 or p53. A. Mutations of ELL selectively abrogate interaction with EAF1. IP-western blotting of the chromatin fraction of HEK293TL cells transiently expressing FLAG-tagged GAL4-ELL proteins with or without S600A/K606T substitutions (SA/KT) along with xAF4 and EAF1h was performed as in Figure 4D. B. Recruitment of exogenously expressed AF4 or EAF1 by ELL mutant proteins. ChIP-qPCR analysis of HEK293TL cells transiently expressing FLAG-tagged GAL4-ELL proteins with or without the SA/KT mutation along with xAF4 and EAF1h was performed as in Figure 4C. C. Effects of the SA/KT mutation on MLL-ELL-mediated leukemic transformation ex vivo. Various MLL-ELL constructs with or without the SA/KT mutation were examined for transformation of myeloid progenitors as in Figure 1A. D. Effects of the SA/KT mutation on MLL-ELL-mediated leukemic transformation in vivo. MLL-ELL or its SA/KT mutant was transduced to c-Kit-positive hematopoietic progenitors and transplanted into syngeneic mice (n = 20). Primary leukemia cells were harvested from the bone marrow and transplanted into secondary recipient mice (n = 20). E. Enhanced interaction of ELL with AEP mediated by the TRX2 domain. IP-western blotting of the chromatin fraction of virus packaging cells transiently expressing various HA-tagged MTMT-ELL fusion constructs (with or without the SA/KT mutation), was performed. Endogenous proteins co-purified with MLL-ELL proteins were visualized by specific antibodies.

Next, we examined the effects of the SA/KT mutation on the transforming properties of MLL-ELL. Both *Hoxa9* expression and colony-forming potentials were maintained by MLL-ELL SA/KT-ICs (Figures 6C and S6C). Remarkably, the MTM construct fused to ELL carrying the SA/KT mutation (see MTMh-ELĹ’ SA/KT) activated *Hoxa9* and immortalized HPCs albeit with low clonogenicity despite the lack of TRX2 domain, suggesting that the SA/KT mutation partially compensates for the lack of interaction with the HBO1 complex. MLL-ELL SA/KT induced leukemia in vivo (Figure 6D), indicating that direct recruitment of EAF1 or p53 is dispensable for MLL-ELL-mediated leukemic transformation. sgRNA competition assays showed that loss of *Eaf1* has a marginal inhibitory effect on proliferation of MLL-ELL-ICs ex vivo, while loss of *Trp53* accelerated it, suggestive of non-essential roles for EAF1 and an inhibitory role for p53 in MLL-ELL-mediated transformation (Figure S6D). IP-western blotting of the MTMh- or MTMTh-ELĹ’ SA/KT showed that association with AF4 and ENL through the presumed secondary contact is enhanced by the presence of TRX2 domain while p53 association remained abolished (Figure 6E). Leukemia cells (LCs) of MLL-ELL were particularly sensitive to WM1119, a pan MYST family HAT inhibitor (MacPherson *et al*., 2020), compared to other MLL fusion-LCs (Figure S6E), suggesting that HBO1-mediated protein acetylation may be implicated in the enhancement of ELL-AF4 association. In summary, the results suggested that MLL-ELL transforms HPCs via interaction with AF4 family proteins, which is normally promoted by the HBO1 complex but hindered by other ELL-associated factors.

### NUP98-HBO1 fusion transforms myeloid progenitors through interaction with MLL

NUP98-HBO1 fusion has been found in chronic myelomonocytic leukemia and was shown to induce clinically relevant leukemia in mice (Hayashi *et al*, 2019). NUP98-HBO1 transformed HPCs ex vivo, accompanying with high level *Hoxa9* expression (Figure 7A). Fusion of NUP98 and homeodomain proteins (e.g., NUP98-HOXA9) has been shown to induce leukemia in mouse models (Kroon *et al*, 2001). The homeodomain of HOXA9 possesses a sequence-specific DNA binding ability, demonstrated by ChIP-qPCR analysis on a HOXA9 binding site within the HOXA7 locus (Figure S7A). A FLAG-tagged NUP98-HOXA9 construct (fNUP98-HOXA9) bound to the HOXA9 binding site, whereas fNUP98-HBO1 did not, suggesting that HBO1 provides an alternative chromatin targeting function which differs from that of HOXA9 homeodomain. Because the HBO1 complex binds MLL (Figure 7B), we hypothesized that the HBO1 portion may confer a targeting ability through association with MLL. To test this hypothesis, we generated an artificial NUP98 fusion construct in which MENIN is fused to NUP98 and examined its transforming properties. Indeed, NUP98-MENIN transformed HPCs (Figure 7C). Artificial NUP98 constructs fused with an HBO complex component (i.e., fNUP98-MEAF6 or -ING5) also transformed HPCs. IP-western blotting showed that these artificial NUP98 fusions associated with MLL in the chromatin fraction, supporting the hypothesis that NUP98-HBO1 transforms HPCs via interaction with MLL (Figure 7D). The 5’ portion of NUP98 did not co-precipitate MLL in this setting (see fNUP98-5’), consistent with a previous report (Shima *et al*, 2017). However, it should be noted that the 5’ portion of NUP98 was shown to localize in proximity with MLL (Xu *et al*, 2016) and promote the physical interaction of NUP98-HOXA9 with MLL (Shima *et al*., 2017). The sgRNA competition assay demonstrated fNUP98-HBO1-ICs depended on *Mll* for continuous proliferation (Figure 7E), similarly to fNUP98-HOXA9-ICs as reported previously (Shima *et al*., 2017; Xu *et al*., 2016). However, both fNUP98-HOXA9- and fNUP98-HBO1-ICs were mildly sensitive to MENIN-MLL interaction inhibitor (MI-2-2) (Shi *et al*, 2012), and were not rendered completely differentiated unlike MLL-AF10-ICs (Figure 7F), suggesting that a MENIN-less MLL complex may sufficiently recruit NUP98-HBO1 to the target chromatin. Interestingly, E508Q amino acid substitution, which kills its HAT activity (Foy *et al*, 2008) but retains the binding capacities to MLL, did not impair the transforming property of NUP98-HBO1 (Figure 7C, D, see fNUP98-HBO1 EQ). Moreover, WM1119 did not induce complete differentiation of NUP98-HBO1-ICs unlike MLL-AF10-ICs (Figure S7B). These results indicate that interaction with MLL, but not the intrinsic HAT activity, mediates NUP98-HBO1-mediated leukemic transformation.

**Figure 7.**
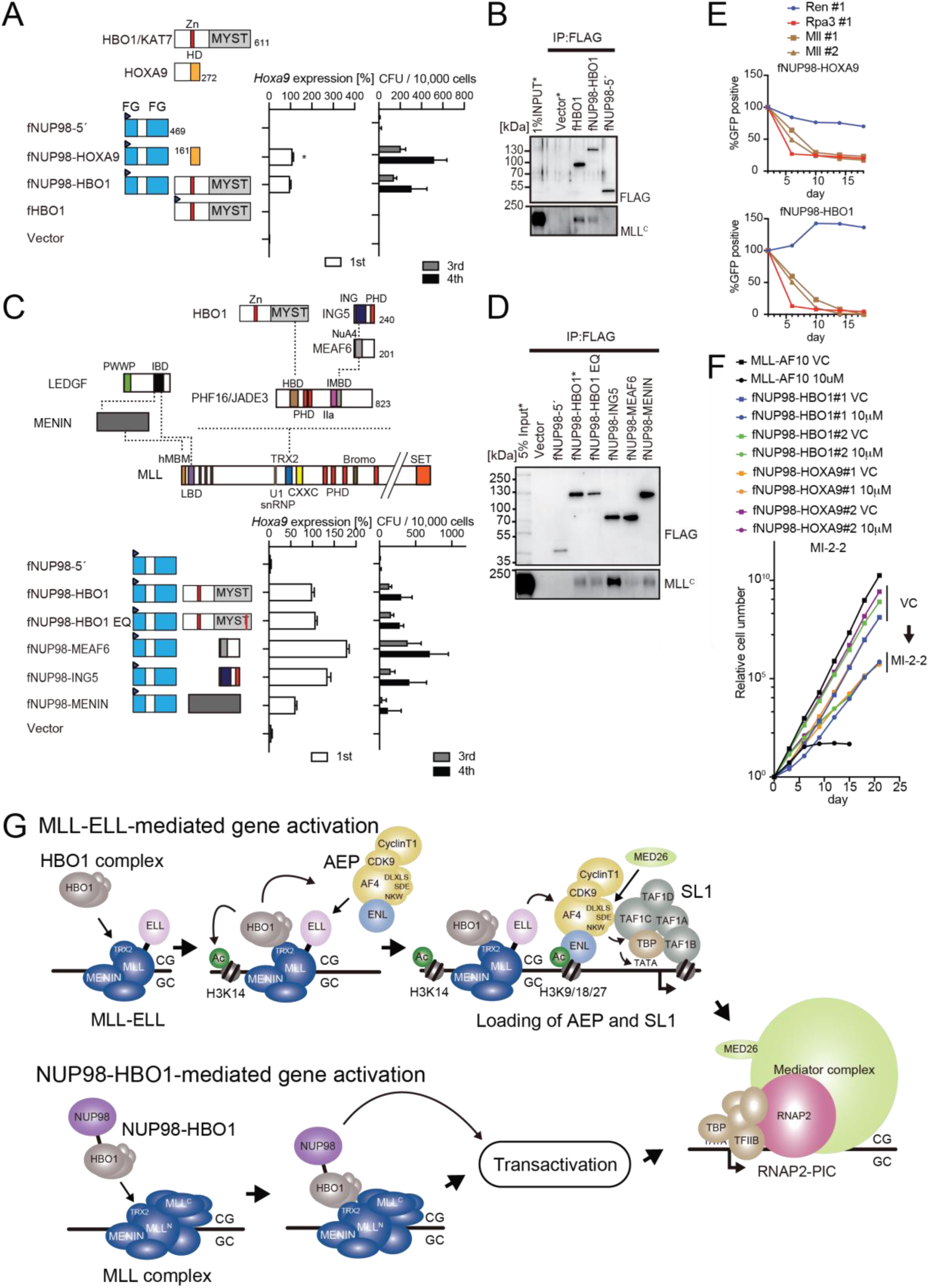
NUP98-HBO1 fusion transforms myeloid progenitors through interaction with MLL. A. Structural requirement of NUP98-HBO1 fusion. Various NUP98 fusion constructs were examined for transformation of myeloid progenitors (along with a HBO1 construct) as in Figure 1A. Relative *Hoxa9* expression in first round colonies was analyzed. Asterisk indicates that the qPCR probe detected human HOXA9 coding sequence included in the NUP98-HOXA9 construct in addition to endogenous murine *Hoxa9*. B. Association with MLL by NUP98-HBO1. The chromatin fraction of virus packaging cells transiently expressing the indicated transgenes was subjected to IP-western blotting. Endogenous MLL^C^ fragment was detected by specific anti-MLL antibody. C. Structural requirement of various NUP98 fusions for leukemic transformation. Indicated NUP98 fusion constructs were examined for transformation of myeloid progenitors as in Figure 1A. D. Association with MLL by various NUP98 fusions. IP-western blotting was performed as in Figure 7B. E. Requirement of the *Mll* gene for NUP98 fusion-immortalized cells. sgRNA competition assays for *Mll* were performed on NUP98-HOXA9- and NUP98-HBO1-immortalized cells as in Figure 2F. F. Effects of pharmacologic inhibition of MENIN-MLL interaction on NUP98 fusion-immortalized cells. NUP98-HBO1- and NUP98-HOXA9-immortalized cells were cultured in the presence of 10 μM of MI-2-2 MENIN-MLL interaction inhibitor, and their proliferation was monitored every 3 days. MLL-AF10-immortalized cells were also analyzed as a positive control. VC: vehicle control G. Models of MLL-ELL- and NUP98-HBO1-mediated gene activation

## Discussion

Most MLL fusion proteins constitutively recruit AEP to immortalize HPCs (Takahashi & Yokoyama, 2020). However, it was unclear how MLL-ELL exerts its oncogenic properties, because an artificial MLL-EAF1 fusion immortalizes HPCs (Luo *et al*., 2001). AF4- and EAF-family proteins share structural similarities, both of which have the SDE motif which binds SL1 (Figure 5) (Okuda *et al*., 2016). We previously demonstrated that the combination of SDE and NKW motifs is required for transcriptional activation (Okuda *et al*., 2015). Because EAF1 does not have an NKW motif, EAF1 is incompetent for transcriptional activation, and therefore MTMh-EAF1-C did not transform HPCs (Figure 5G). Instead, EAF1 homodimerizes through its EHD. The MTMT-EHD fusion (e.g. MTMTh-EAF1-N) transformed HPCs in TRX2 domain-dependent manner, similar to MLL-AF6, which is known to transform HPCs via homodimerization (Liedtke *et al*., 2010). Thus, we concluded that MLL-EAF1 transforms HPCs via homodimerization. The S600A/K606T double mutation on ELL completely abrogated the interaction with EAF1, while retaining some binding capacity to AF4, and did not impair MLL-ELL-mediated transformation. Thus, we concluded that MLL-ELL transforms HPCs via association with AF4 family proteins, but not with EAF1 (Figure 7G).

ChIP-qPCR analysis indicated that ELL recruited AF4 and ENL to the target chromatin, but was unable to further recruit SL1 and MED26 (Figure 5D), suggesting that the AEP complex must dissociate from ELL to function as a transcriptional activator. Thus, MLL-ELL is a loading factor, but not a nucleating factor (like MLL-ENL), of the AEP complex. In our previous study, we showed that MLL-AF10 is also an AEP loading factor which loads ENL onto chromatin (Okuda *et al*., 2017). It remains unclear how MLL-AF6, a dimer type of MLL fusion, affects the function of AEP and transforms HPCs. Nonetheless, ChIP-qPCR analysis showed that MLL-AF6 co-localized with AEP components at its target promoters (Yokoyama *et al*., 2010), suggesting that MLL-AF6 also loads AEP to the target chromatin via unknown mechanisms. All of the presumed AEP loading factor type MLL fusions (i.e. MLL-ELL, MLL-AF10, and MLL-AF6) showed susceptibility to *Enl* knockdown, while the AEP nucleating factor type MLL fusions (e.g., MLL-ENL) were relatively resistant (Figure S5B) (Okuda *et al*., 2017; Yokoyama *et al*., 2010). Moreover, the AEP loading factor type MLL fusions exhibited some degree of dependency on the TRX2 domain, which mediates HBO1 complex recruitment (Figures 1 and S1). These results suggest that the HBO1 complex promotes loading of the AEP complex onto target chromatin (Figure 7G). Hence, the inhibitors that interfere with the MLL-HBO complex interaction may be highly effective for the AEP loading factor type MLL fusions.

Like MLL, NUP98 fuses with a variety of partners (Gough *et al*, 2011). NUP98 fusion partners can be roughly subdivided into three groups; the homeodomain containing protein type (e.g., NUP98-HOXA9), the chromatin reader type (e.g., NUP98-HBO1, - LEDGF, -MLL, -NSD1, -PHF23, -BPTF, and -KDM5A), and the coiled-coil structure containing protein type. Among the chromatin reader type NUP98 fusions, MLL and LEDGF are components of the MLL complex (Yokoyama & Cleary, 2008), which targets previously transcribed CpG-rich promoters through association with unmethylated CpGs and di/tri-methylated histone H3 lysine 36 (H3K36me2/3) (Okuda *et al*., 2014), and di/tri-methylated Histone H3 lysine 4 (H3K4me2/3) (Milne *et al*, 2010; Wang *et al*, 2010). NUP98-NSD1 was shown to activate HOX-A genes through the association with methylated H3K36 marks (Wang *et al*, 2007), and NUP98-PHF23, -BPTF, and-KDM5A were through association with H3K4me2/3 marks (Zhang *et al*, 2020). Thus, the chromatin reader type NUP98 fusions confer a targeting ability to the chromatin of the HOX-A loci by interacting with specific chromatin modifications and/or chromatin binding proteins. Some NUP98 fusions including NUP98-HBO1 and -LEDGF appear to target through interaction with MLL. This was evident from the immortalization of HPCs by the artificial fusion of NUP98 and MENIN (Figure 7C). It has been reported that a subset of MLL target genes are regulated in a MENIN-independent manner (Artinger *et al*, 2013). In addition, an artificial construct of MLL, lacking the MENIN-binding motif, but retaining the CXXC domain and the PHD finger 3 localized at the *HOXA9* locus, presumably through the association with unmethylated CpGs and H3K4me2/3(Milne *et al*., 2010; Wang *et al*., 2010), suggesting that wild type MLL can bind to some of its target chromatin in a MENIN-independent manner. The NUP98-MLL fusion genes found in patients do not retain the structures responsible for interaction with MENIN and LEDGF (Kaltenbach *et al*, 2010), suggesting that the MLL portion in NUP98-MLL confers its targeting ability in a MENIN-independent manner. NUP98-HBO1-ICs proliferate depending on MLL, but are relatively resistant to the MENIN-MLL interaction inhibitor compared with MLL-AF10-ICs (Figure 7), indicating that a MENIN-less MLL complex may sufficiently recruit NUP98-HBO1. Taken together, the results suggest that a subset of NUP98 fusions target the HOX-A loci via association with MLL and activate transcription by NUP98-mediated functions (Figure 7G).

This study demonstrated that various MLL fusions and NUP98 fusions transform HPCs via HBO1-MLL interaction (Figure 7G). Hence, we propose that the binding modules for this interaction can be a good molecular target for drug development. HBO1-MLL interaction inhibitors would complement the therapeutic effects of the emerging MENIN-MLL interaction inhibitors (Klossowski *et al*, 2019; Krivtsov *et al*, 2019) to treat malignant leukemia with subsets of MLL- and NUP98-fusions.

## Materials and Methods

### Materials Availability

Materials generated in this study will be provided upon request.

### Data and Code availability

ChIP-seq data, CIRA-seq data, and RNA-seq data have been deposited at the DDBJ (DNA Data Bank of Japan) Sequence Read Archive under the accession numbers (DRA010818, DRA004871, DRA004872, DRA010819, DRA008732, DRA008734, and, DRA004874) and sample IDs listed in Tables S4.

## Experimental model and subject details

### Vector constructs

For protein expression vectors, cDNAs obtained from Kazusa Genome Technologies Inc. (Nagase *et al*, 2008) were modified by PCR-mediated mutagenesis and cloned into the pMSCV vector (for virus production) or pCMV5 vector (for transient expression) by restriction enzyme digestion and DNA ligation. The MSCV-neo MLL-ENL, and MLL-AF10 vectors have been previously described (Okuda *et al*., 2017). sgRNA-expression vectors were constructed using the pLKO5.sgRNA.EFS.GFP vector (Heckl *et al*, 2014). shRNA-expression vectors were constructed using a pLKO.1 vector, or were purchased from Dharmacon. The target sequences are listed in Table S2 (sgRNA).

### Cell lines

HEK293T cells were a gift from Michael Cleary and were authenticated by the JCRB Cell Bank in 2019. HEK293TN cells were purchased from System Biosciences. The cells were cultured in Dulbecco’s modified Eagle’s medium (DMEM), supplemented with 10% fetal bovine serum (FBS) and penicillin-streptomycin (PS). The Platinum-E (PLAT-E) ecotropic virus packaging cell line—a gift from Toshio Kitamura (Morita *et al*, 2000)—was cultured in DMEM supplemented with 10% FBS, puromycin, blasticidin, and PS. The human leukemia cell lines HB1119 and REH, gifts from Michael Cleary (Tkachuk *et al*, 1992), was cultured in RPMI 1640 medium supplemented with 10% FBS and PS. Cells were incubated at 37 °C in a 5% CO_2_ atmosphere. HEK293T–LUC cells were generated by transduction of the lentivirus carrying pLKO1-puro-FR-LUC, as described previously (Okuda *et al*., 2015). Murine myeloid progenitors immortalized by various transgenes were cultured in RPMI 1640 medium supplemented with 10% FBS, and PS containing murine stem cell factors, interleukin-3, and granulocyte-macrophage colony-stimulating factor (1 ng/mL of each).

### Western blotting

Western blotting was performed as previously described (Yokoyama *et al*, 2002). Briefly, proteins were separated electrophoretically in an acrylamide gel and were transblotted onto nitrocellulose sheets using a mini transblot cell (Bio-Rad). The nitrocellulose sheets were blocked with 5% skimmed milk in T-PBS (phosphate-buffered saline containing 0.1% Tween 20) for 1 h, rinsed twice with T-PBS, and incubated with primary antibodies suspended in 5% skim milk in T-PBS overnight. The blots were then washed twice with T-PBS and incubated with peroxidase-conjugated secondary antibodies for 2 h. Chemiluminescence was performed using the ECL chemiluminescence reagent (GE Healthcare). The antibodies used in this study are listed in Table S1.

### Virus production

Ecotropic retrovirus was produced using PLAT-E packaging cells (Morita *et al*., 2000). Lentiviruses were produced in HEK293TN cells using the pMDLg/pRRE, pRSV-rev, and pMD2.G vectors, all of which were gifts from Didier Trono (Dull *et al*, 1998). The virus-containing medium was harvested 24–48 h following transfection and used for viral transduction.

### Myeloid progenitor transformation assay

The myeloid progenitor transformation assay was carried out as previously described (Lavau *et al*., 1997; Okuda & Yokoyama, 2017b). Bone marrow cells were harvested from the femurs and tibiae of 5-week-old female C57BL/6J mice. c-Kit^+^ cells were enriched using magnetic beads conjugated with an anti-c-Kit antibody (Miltenyi Biotec), transduced with a recombinant retrovirus by spinoculation, and then plated (4 × 10^4^ cells/sample) in a methylcellulose medium (Iscove’s modified Dulbecco’s medium, 20% FBS, 1.6% methylcellulose, and 100 µM β-mercaptoethanol) containing murine stem cell factor (mSCF), interleukin 3 (mIL-3), and granulocyte-macrophage colony-stimulating factor (mGM-CSF; 10 ng/mL each). During the first culture passage, G418 (1 mg/mL) or puromycin (1 µg/mL) was added to the culture medium to select the transduced cells. *Hoxa9* expression was quantified by RT-qPCR after the first passage. Cells were then re-plated once every 5 days with fresh medium. Colony-forming units were quantified per 10^4^ plated cells at each passage.

### In vivo leukemogenesis assay

In vivo leukemogenesis assay was carried out as previously described (Lavau *et al*., 1997; Okuda & Yokoyama, 2017a). Cells positive for c-Kit (2 × 10^5^), prepared from mouse femurs and tibiae, were transduced with retrovirus by spinoculation and intravenously transplanted into sublethally irradiated (5-6 Gy) 8-week-old female C57BL/6JJcl (C57BL/6J) mice. Moribund mice were euthanized and the cells isolated from their bone marrow were cultured in methylcellulose medium used for myeloid progenitor transformation assays for more than three passages to remove untransformed cells and then subjected to secondary transplantation. For secondary leukemia, leukemia cells (2 × 10^5^) cultured ex vivo were transplanted in the same manner as the primary transplantation. This protocol was approved by the National Cancer Center Institutional Animal Care and Use Committee.

### qRT-PCR

Total RNA was isolated using the RNeasy Mini Kit (Qiagen) and reverse-transcribed using the Superscript III First Strand cDNA Synthesis System (Thermo Fisher Scientific) with oligo (dT) primers. Gene expression was analyzed by qPCR using TaqMan probes (Thermo Fisher Scientific). Relative expression levels were normalized to those of *GAPDH*/*Gapdh* and determined using a standard curve and the relative quantification method, according to the manufacturer’s instructions (Thermo Fisher Scientific). Commercially available PCR probe sets were used are listed in Table S1.

### Fractionation-assisted chromatin immunoprecipitation (fanChIP)

Chromatin fractions from HEK293T and HB1119 cells were prepared using the fanChIP method, as described previously (Okuda *et al*., 2014). Cells were suspended in CSK buffer (100 mM NaCl, 10 mM PIPES [pH 6.8], 3 mM MgCl_2_, 1 mM EGTA, 0.3 M sucrose, 0.5% Triton X-100, 5 mM sodium butyrate, 0.5 mM DTT, and protease inhibitor cocktail) and centrifuged (400 × *g* for 5 min, at 4 °C) to remove the soluble fraction. The pellet was resuspended in MNase buffer (50 mM Tris-HCl [pH 7.5], 4 mM MgCl_2_, 1 mM CaCl_2_, 0.3 M sucrose, 5 mM sodium butyrate, 0.5 mM DTT, and protease inhibitor cocktail) and treated with MNase at 37 °C for 3–6 min to obtain oligonucloesomes. MNase reaction was then stopped by adding EDTA (pH 8.0) to a final concentration of 20 mM. An equal amount of lysis buffer (250 mM NaCl, 20 mM sodium phosphate [pH 7.0], 30 mM sodium pyrophosphate, 5 mM EDTA, 10 mM NaF, 0.1% NP-40, 10% glycerol, 1 mM DTT, and EDTA-free protease inhibitor cocktail) was added to increase solubility. The chromatin fraction was cleared by centrifugation (15000rpm for 5m, 4 °C) and subjected to immunoprecipitation with specific antibodies and Protein-G magnetic microbeads (Invitrogen) or with anti-FLAG M2 antibody-conjugated beads (Table S1). Immunoprecipitates were then washed five times with washing buffer (1:1 mixture of lysis buffer and MNase buffer with 20 mM EDTA) and eluted in elution buffer (1% SDS and 50 mM NaHCO_3_). The eluted material was analyzed by multiple methods including western blotting, qPCR, and deep sequencing. Optionally, the immunoprecipitates were washed twice with washing buffer, twice with MNase buffer, and then treated with DNase I (Qiagen) for 15 min with room temperature. The immunoprecipitates were further washed four times with washing buffer, and eluted using elution buffer. The eluted materials were subjected to western blotting and SYBR green staining.

### ChIP-qPCR and ChIP-seq

The eluted material obtained by fanChIP was extracted using phenol/chloroform/isoamyl alcohol. DNA was precipitated with glycogen, dissolved in TE buffer, and analyzed by qPCR (ChIP-qPCR) or deep sequencing (ChIP-seq). The qPCR probe/primer sequences are listed in Table S3. For deep sequencing, DNA was further fragmented (∼150 bp) using the Covaris M220 DNA shearing system (M&M Instruments Inc.). Deep sequencing was then performed using the TruSeq ChIP Sample Prep Kit (Illumina) and HiSeq2500 (Illumina) at the core facility of Hiroshima University. Data were visualized using Integrative Genome Viewer (The Broad Institute). Raw reads in FASTQ format were trimmed using Cutadapt and aligned to the reference genome hg19 with BWA (Li & Durbin, 2009; Martin, 2011). Accession numbers and sample IDs are listed in the Table S4.

### Liquid chromatography-tandem mass spectrometry (LC-MS/MS) analysis

Proteins were digested with trypsin and tandem mass spectrometry was performed using an LTQ Orbitrap ELITE ETD mass spectrometer (Thermo Fisher Scientific), as described previously (Okuda *et al*., 2017).

### sgRNA competition assay

*Cas9* was introduced via lentiviral transduction using the pKLV2-EF1a-Cas9Bsd-W vector (Tzelepis *et al*, 2016). Cas9-expressing stable lines were established with blasticidin (10–30 µg/mL) selection. The sequences of sgRNAs are listed in Table S2. The targeting sgRNA was co-expressed with GFP via lentiviral transduction using pLKO5.sgRNA.EFS.GFP vector (Heckl *et al*., 2014). Percentages of GFP^+^ cells were initially determined by FACS analysis at 2 days after sgRNA transduction, and then measured every 4 days.

### Accession numbers

Deep sequencing data used in this study have been deposited in the DNA Data Bank of Japan (DDBJ) Sequence Read Archive under the accession numbers listed in Table S3 (ChIP-seq, CIRA-seq, and RNA-seq)

### Statistical analysis

Statistical analysis was performed using GraphPad Prism 7 software. Data are presented as the mean with standard deviation (SD). Multiple comparisons were performed by two-way ANOVA. The statistical tests were two-sided. Mice transplantation experiments were analyzed by the log-rank test and Bonferroni correction was applied for multiple comparisons. *P* values < 0.05 were considered statistically significant. n.s.: *P* > 0.05, *: *P* ≤ 0.05, **: *P* ≤ 0.01, ***: *P* ≤ 0.001, and ****: *P* ≤ 0.0001.

## Acknowledgments

We thank Makiko Okuda, Yuzo Sato, Megumi Nakamura, Etsuko Kanai, Aya Nakayama and Ayako Yokoyama for technical assistance. We also thank the Shonai Regional Industry Promotion Center members for their administrative support. This work was supported by the Japan Society for the Promotion of Science (JSPS) KAKENHI grants (16H05337, 19H03694 to AY). This work was also supported in part by research funds from the Yamagata prefectural government, the City of Tsuruoka, Dainippon Sumitomo Pharma Co. Ltd., and the Friends of Leukemia Research Fund.

## Author contributions

S.T., H.O., R.M., and A.Y. performed experiments; A.K., H.M., and T.I. performed deep sequencing; A.K. analyzed deep sequencing data; T.K. performed mass spectrometry analysis; A.Y. conceived of the project; A.T.K. and A.Y. supervised the project; S.T. and A.Y. co-wrote the paper.

## Supplementary information

**Figure S1.**
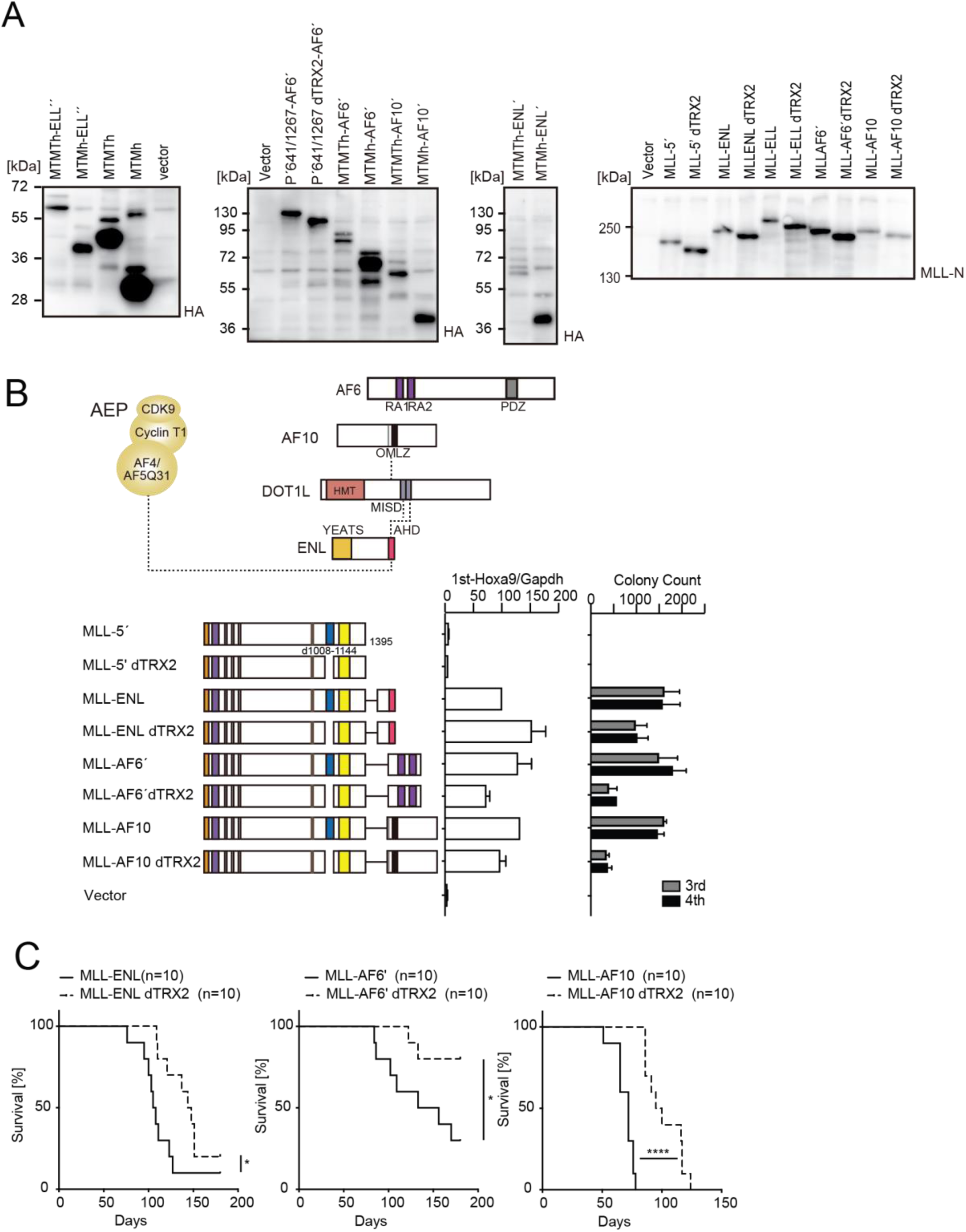
TRX2 domain promotes *MLL*-rearranged leukemogenesis, related to Figure 1. A. Protein expression from various MLL fusion constructs. Virus-packaging cells transiently expressing each MLL fusion construct shown in Figures 1 and S1 were analyzed by western blotting with the indicated antibodies. B. Negative effects imposed by the loss of TRX2 domain on transformation ex vivo. Various MLL fusion constructs lacking the TRX2 domains were examined for transformation of myeloid progenitors as in Figure 1A. C. Negative effects imposed by the loss of TRX2 domain on transformation in vivo. Leukemogenic activity of MLL fusion constructs was examined. Each MLL fusion construct was transduced to c-Kit-positive hematopoietic progenitors and transplanted into syngeneic mice (n=10). Statistical analysis was performed using the log-rank test and Bonferroni correction with the wild type control. \**P* ≤ 0.05. ****: *P* ≤ 0.0001.

**Figure S2.**
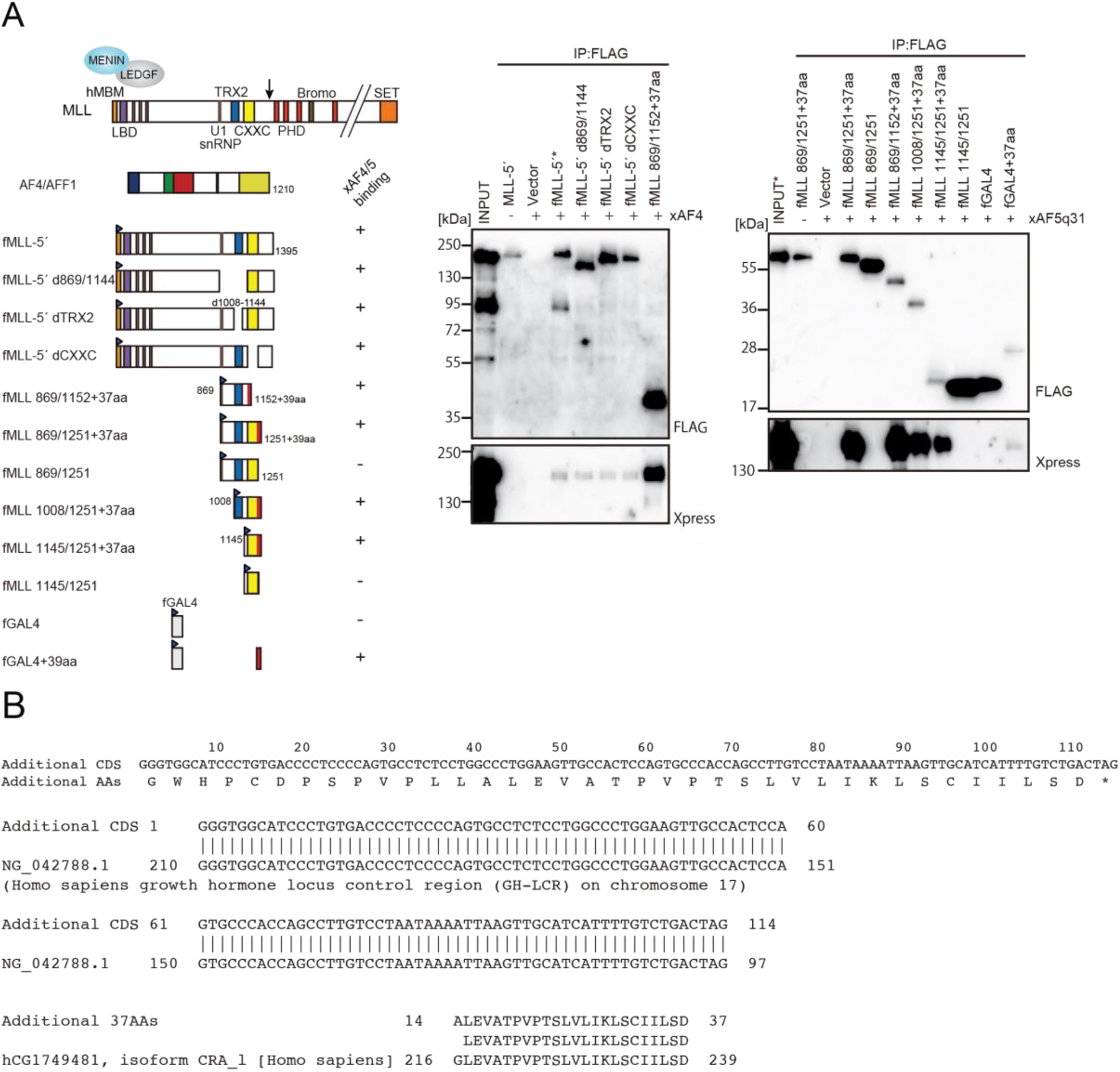
TRX2 domain does not mediate association with AF4 family proteins, related to Figure 2. A. AF4 family proteins bind the additional 37 residues derived from the expression vector. Immunoprecipitation-western blotting of the chromatin fraction of HEK293T cells transiently expressing various FLAG-tagged MLL constructs along with xAF4/xAF5Q31, was performed using fanChIP method. B. Sequence of the additional residues derived from the pCMV5 vector. The coding sequence for the additional 37 amino acids were tethered to MLL in frame. A part of it corresponded to the Chorionic somatomammotropin hormone 1 gene.

**Figure S3.**
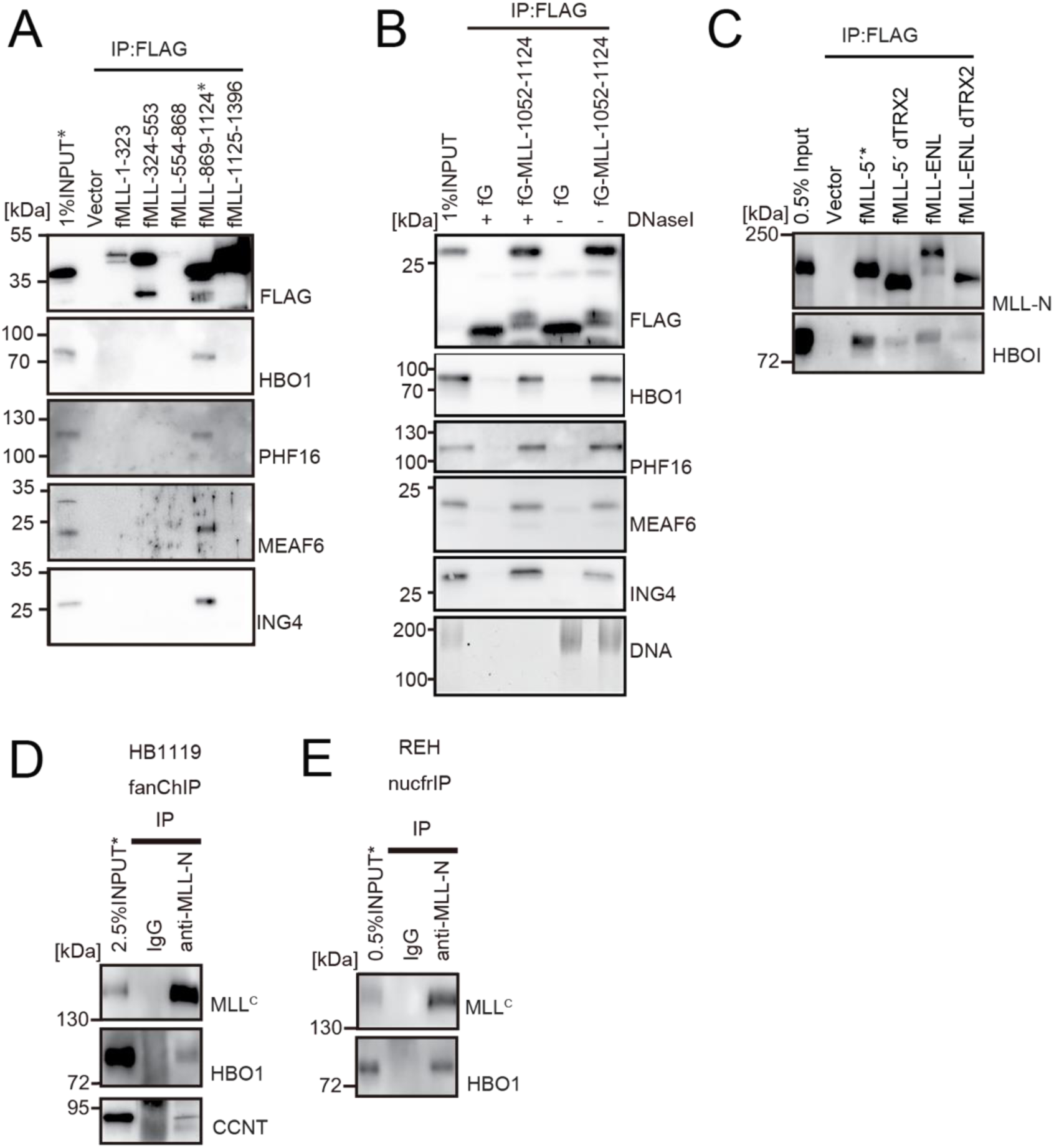
HBO1 complex binds to the TRX2 domain of MLL independently of DNA, related to Figure 2. A. Domain mapping of MLL for interaction with the HBO1 complex. IP-western blotting of the chromatin fraction of HEK293T cells transiently expressing various FLAG-tagged MLL constructs was performed as in Figure 2B. B. Effects of DNase treatment on the interaction between MLL and the HBO complex. Co-precipitates were treated with DNaseI, washed multiple times, and analyzed by western blotting. DNA was detected by SYBR green. C. Interaction with HBO1 through multiple contacts. IP-western blotting of the chromatin fraction of HEK293T cells transiently expressing various FLAG-tagged MLL constructs was performed as in Figure 2B. D. Interaction between MLL proteins and HBO1 in HB1119 leukemia cells. IP-western blotting of the chromatin fraction of HB1119 cells was performed using anti-MLL antibody. E. Interaction between MLL and HBO1 in REH leukemia cells. IP-western blotting of the nucleosome fraction of REH leukemia cells was performed (using anti-MLL antibody) by the nucfrIP method.

**Figure S4.**
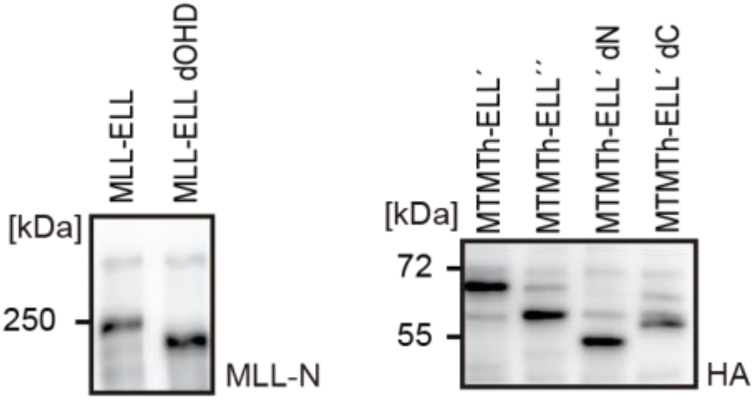
Protein expression from various MLL fusion constructs, related to Figure 4. Virus-packaging cells transiently expressing each MLL fusion construct were analyzed by western blotting with indicated antibodies.

**Figure S5.**
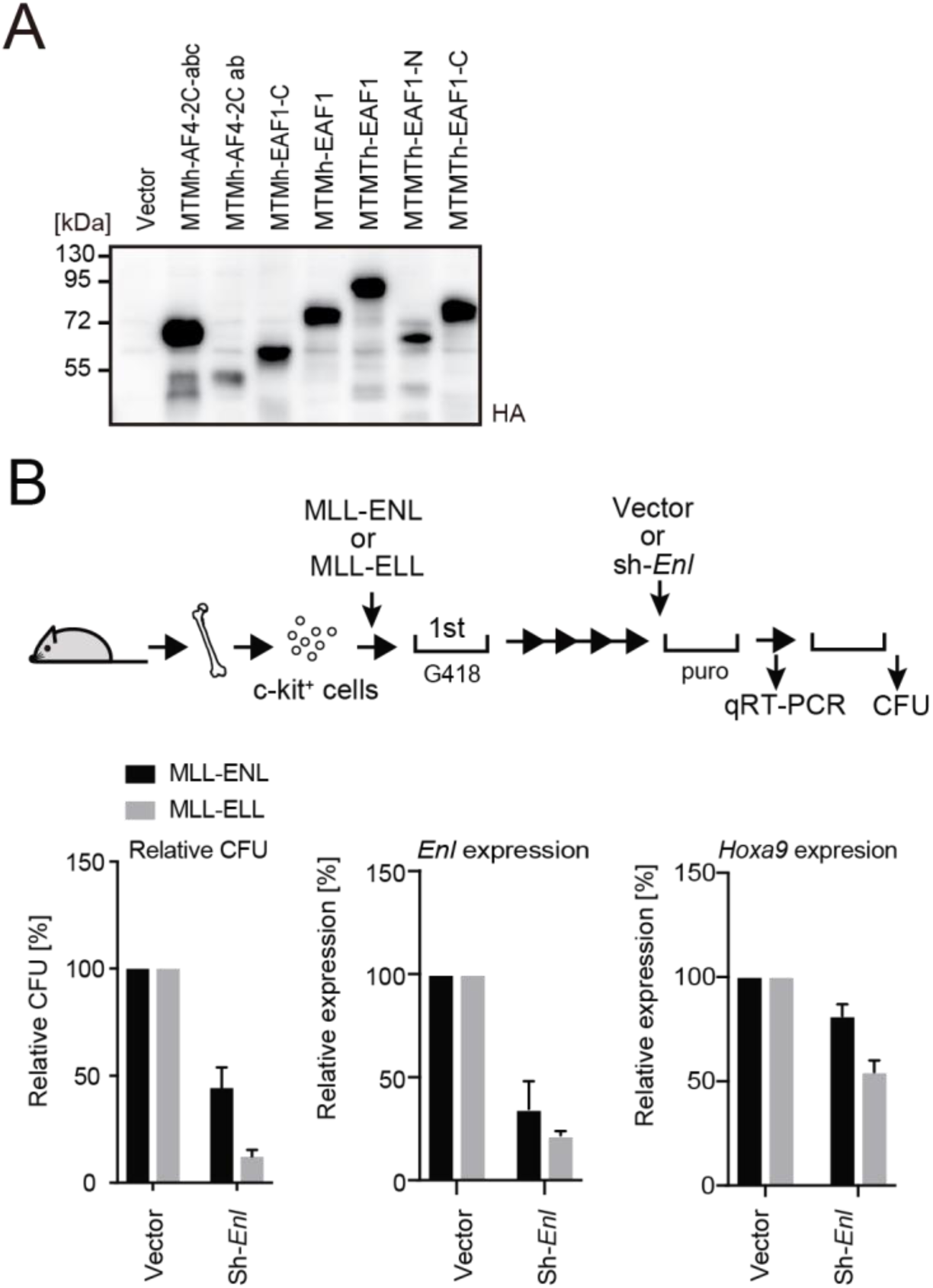
Roles for AEP in MLL-ELL-mediated leukemic transformation, related to Figure 5. A. Protein expression from various MLL fusion constructs. Virus-packaging cells transiently expressing each MLL fusion construct were analyzed by western blotting with indicated antibodies. B. Effects of *Enl* knockdown on MLL-ELL- and MLL-ENL-ICs. MLL-ELL- and MLL-ENL-ICs were transduced with shRNA for endogenous *Enl* and analyzed for gene expression normalized to *Gapdh* by qRT-PCR (mean ± SD of PCR triplicates) and relative colony forming units (mean ± SD of ≥3 biological replicates).

**Figure S6.**
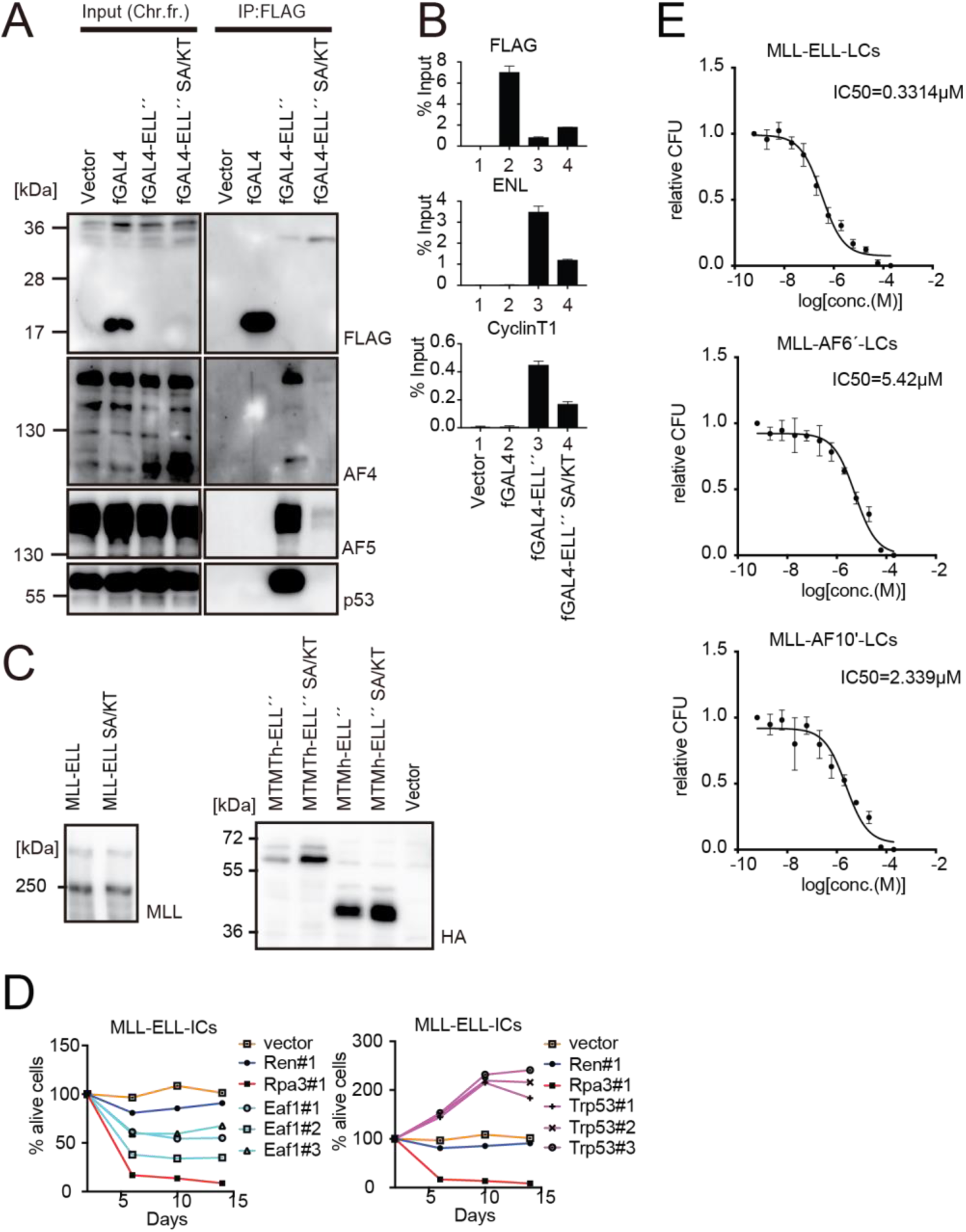
Roles for EAF1 and p53 in MLL-ELL-mediated transformation, related to Figure 6. A. Mutations in ELL selectively abrogated the interaction with p53. IP-western blotting of the chromatin fraction of HEK293TL cells transiently expressing FLAG-tagged GAL4-ELL proteins with or without S600A/K606T substitutions (SA/KT) was performed as in Figure 4D. Endogenous AF4 family proteins and p53 co-purified with GAL4-ELL proteins were detected by anti-AF4-, AF5Q31-, and p53-antibodies. B. Recruitment of endogenous AEP proteins by ELL mutant proteins. ChIP-qPCR analysis of HEK293TL cells transiently expressing FLAG-tagged GAL4-ELL proteins with or without the SA/KT mutation was performed as in Figure 4C. ChIP signals of endogenous ENL and Cyclin T1 were detected using the anti-ENL- and Cyclin T1 antibodies. C. Protein expression from various MLL fusion constructs. Virus-packaging cells, transiently expressing each MLL fusion construct, were analyzed by western blotting with the indicated antibodies. D. Requirement of EAF1 and p53 for MLL-ELL-immortalized cells. sgRNA competition assays for *Eaf1* and *Tp53* were performed as in Figure 2F. E. Sensitivity to pan MYST HAT inhibitor. LCs for various MLL fusions were cultured in the presence of WM1119, a pan MYST HAT inhibitor, and their relative Colony-forming units (mean ± SD of 3 biological replicates) were monitored.

**Figure S7.**
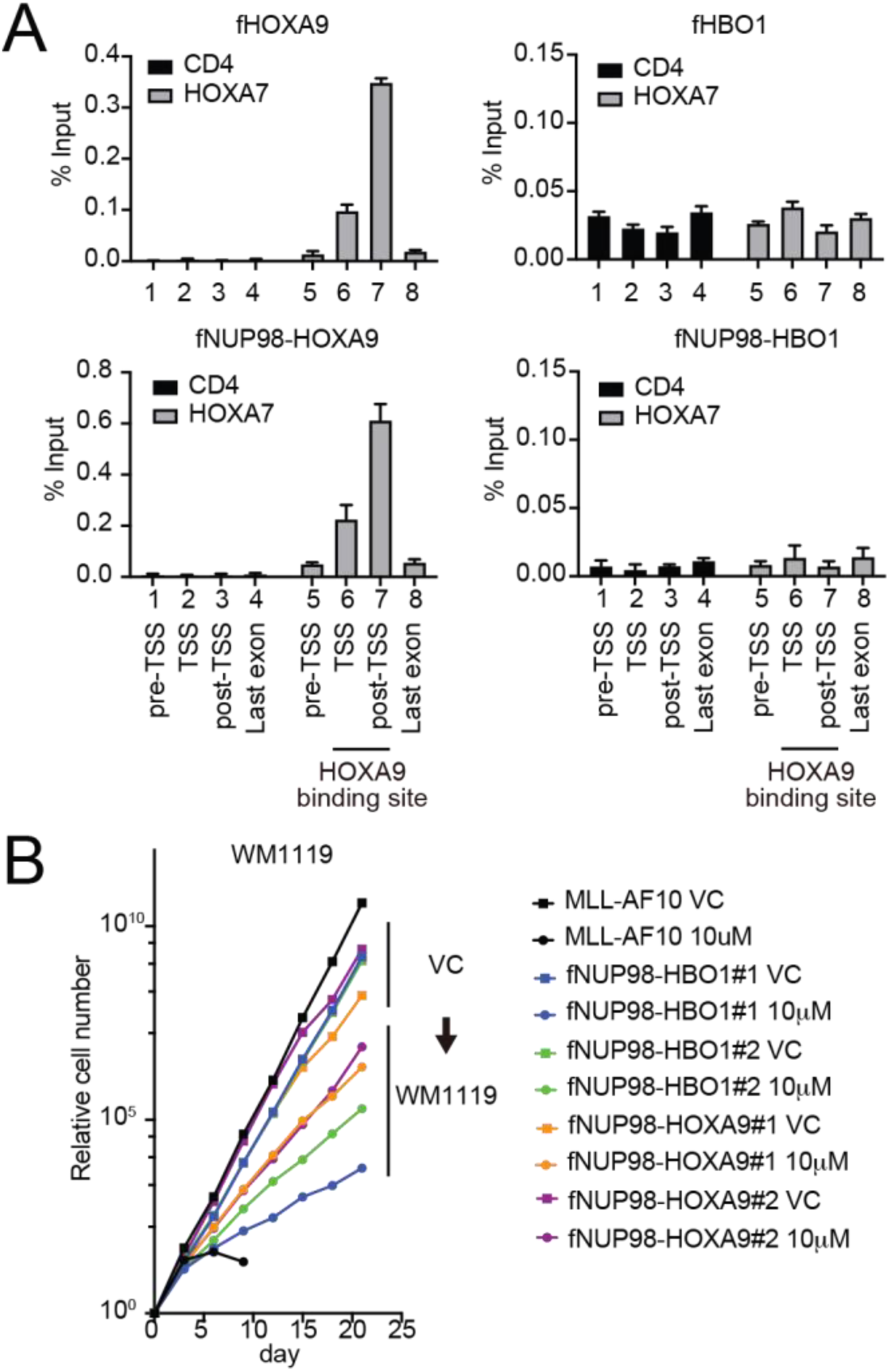
Functional differences between NUP98-HOXA9 and NUP98-HBO1, related to Figure 7. A. Targeting ability to the HOXA9 binding site. Virus packaging cells transiently expressing indicated transgenes were subjected to ChIP-qPCR analysis. The per cent input values of each probe (mean ± SD of PCR triplicates) are shown. A HOXA9 binding site was previously identified at the HOXA7 locus (Shima *et al*, 2017). The CD4 locus was analyzed as a negative control. B. Effects of pharmacologic inhibition of MYST HATs on NUP98 fusion-immortalized cells. NUP98-HBO1- and NUP98-HOXA9-ICs were cultured in the presence of 10 μM of WM1119 MYST HAT inhibitor and their proliferation was monitored every 3 days. MLL-AF10-immortalized cells were also analyzed as positive control. VC: vehicle control

**Table S1.**
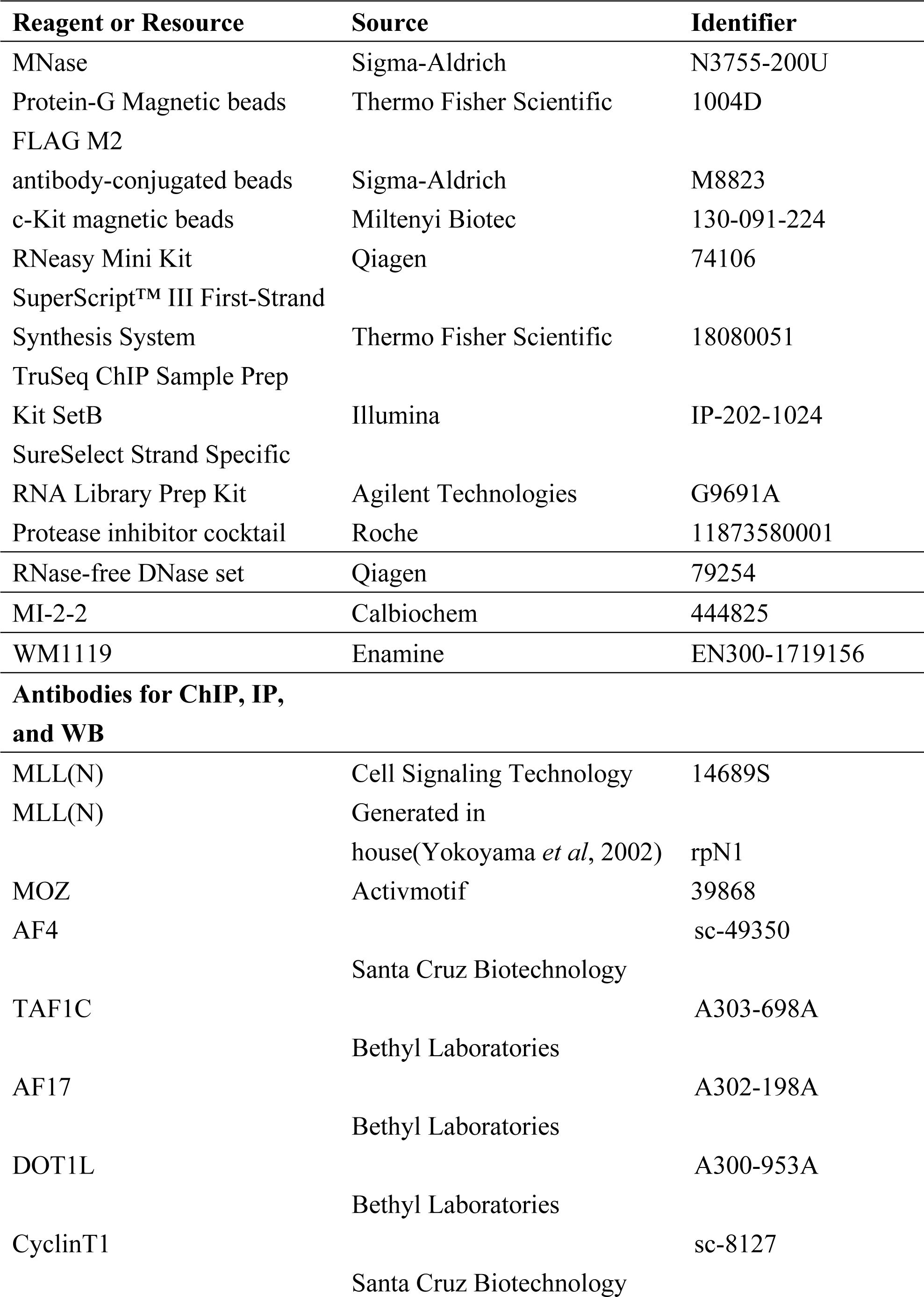
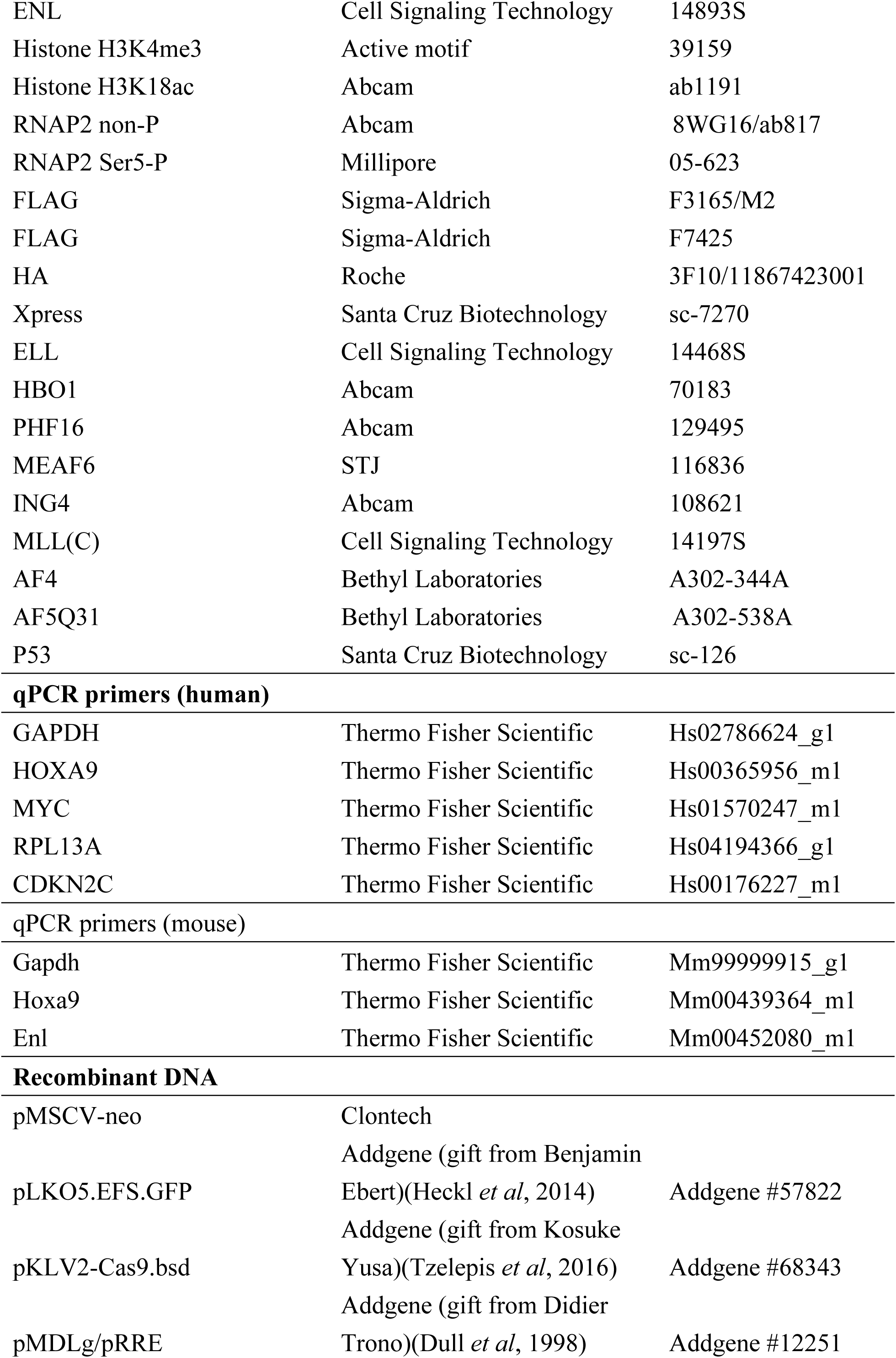
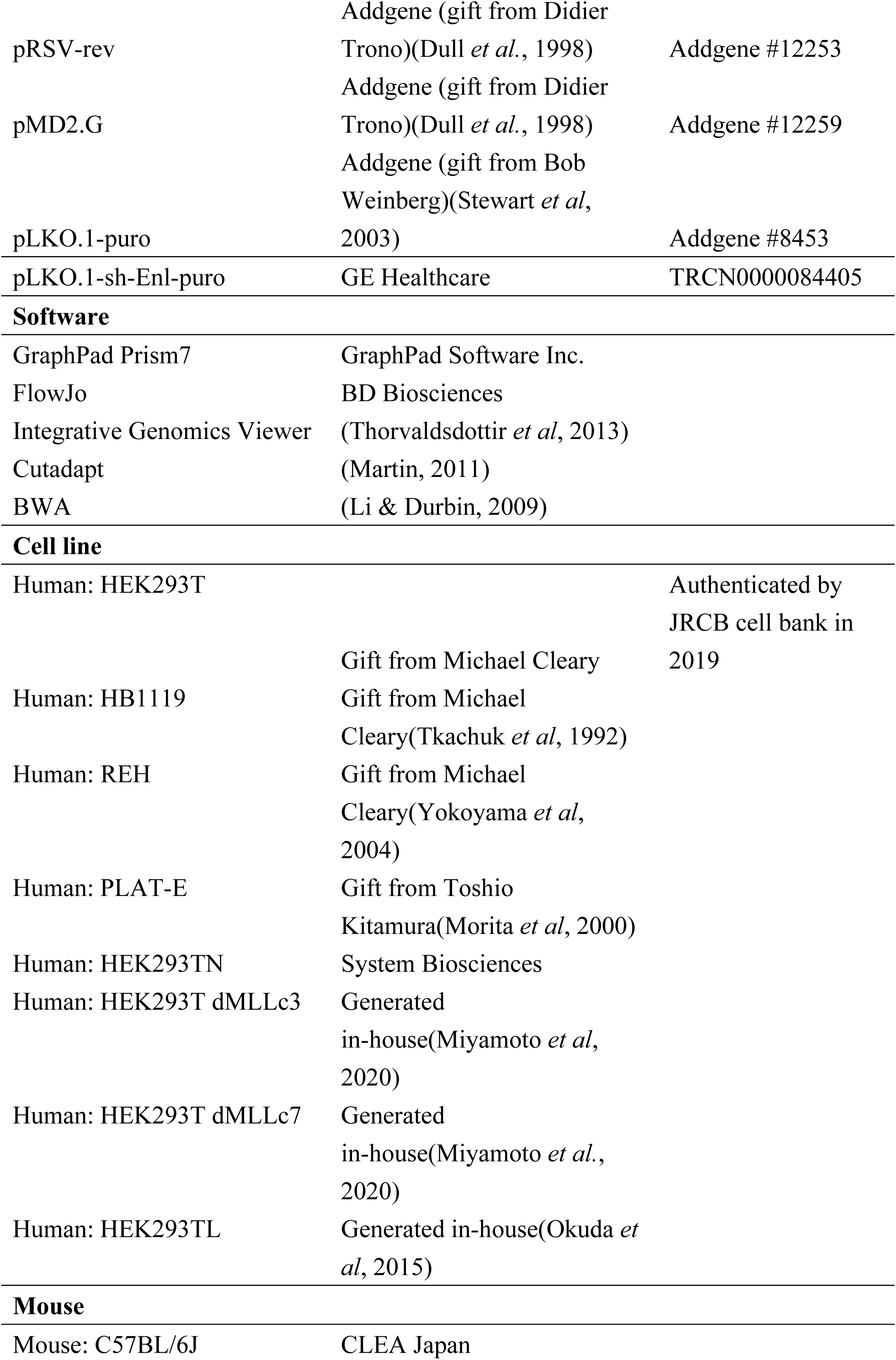
Materials.

**Table S2.**
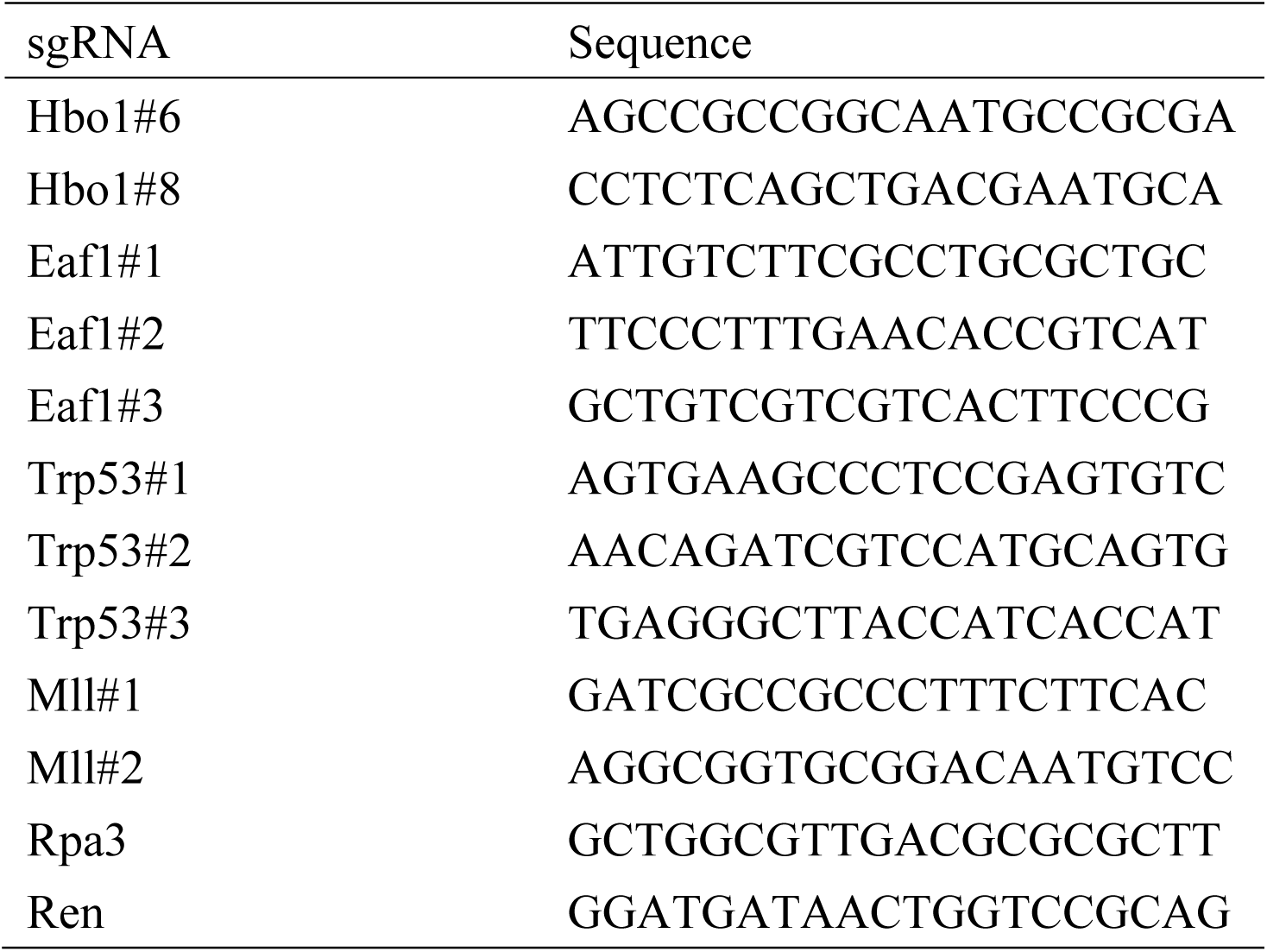
sgRNA sequences.

**Table S3.**
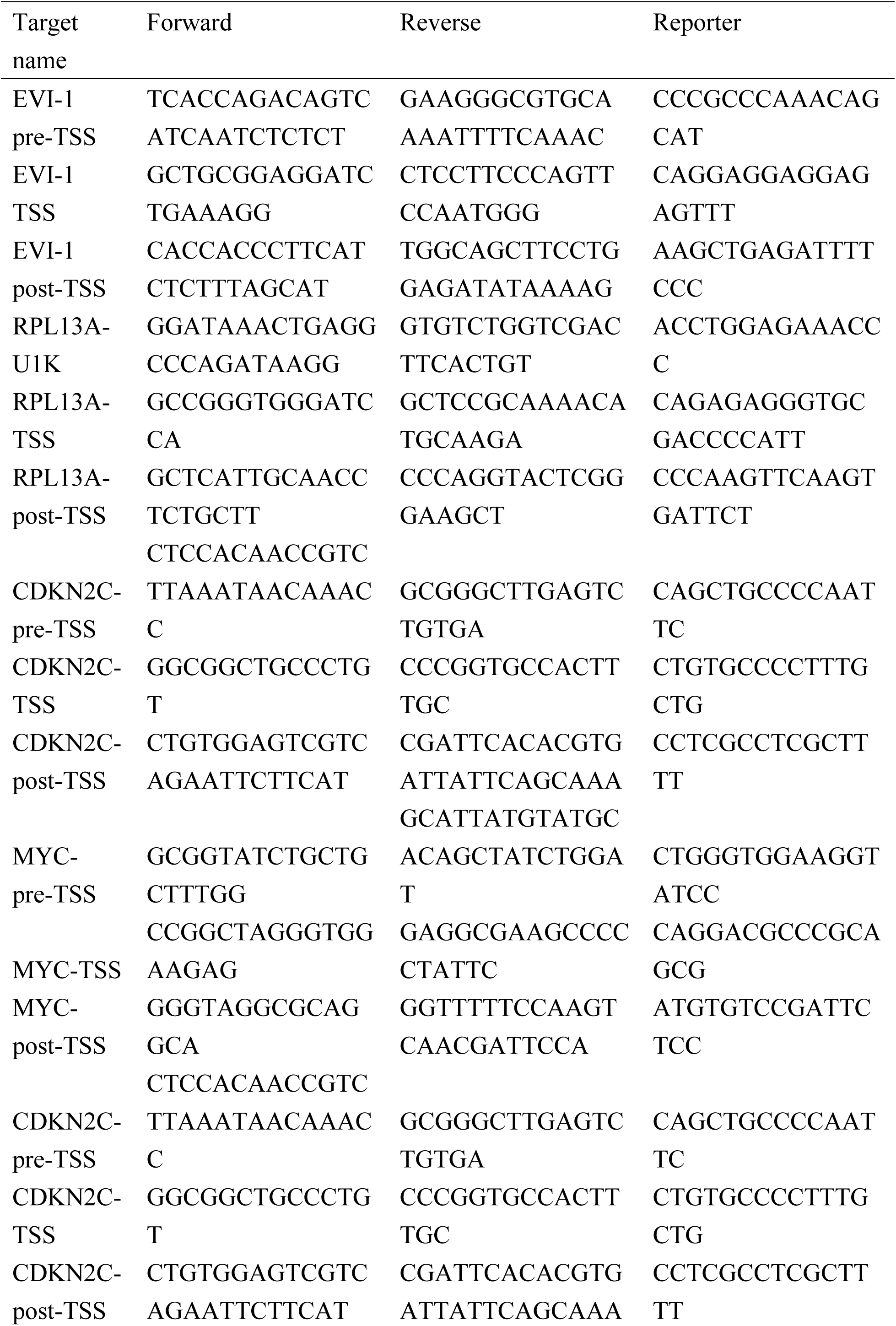
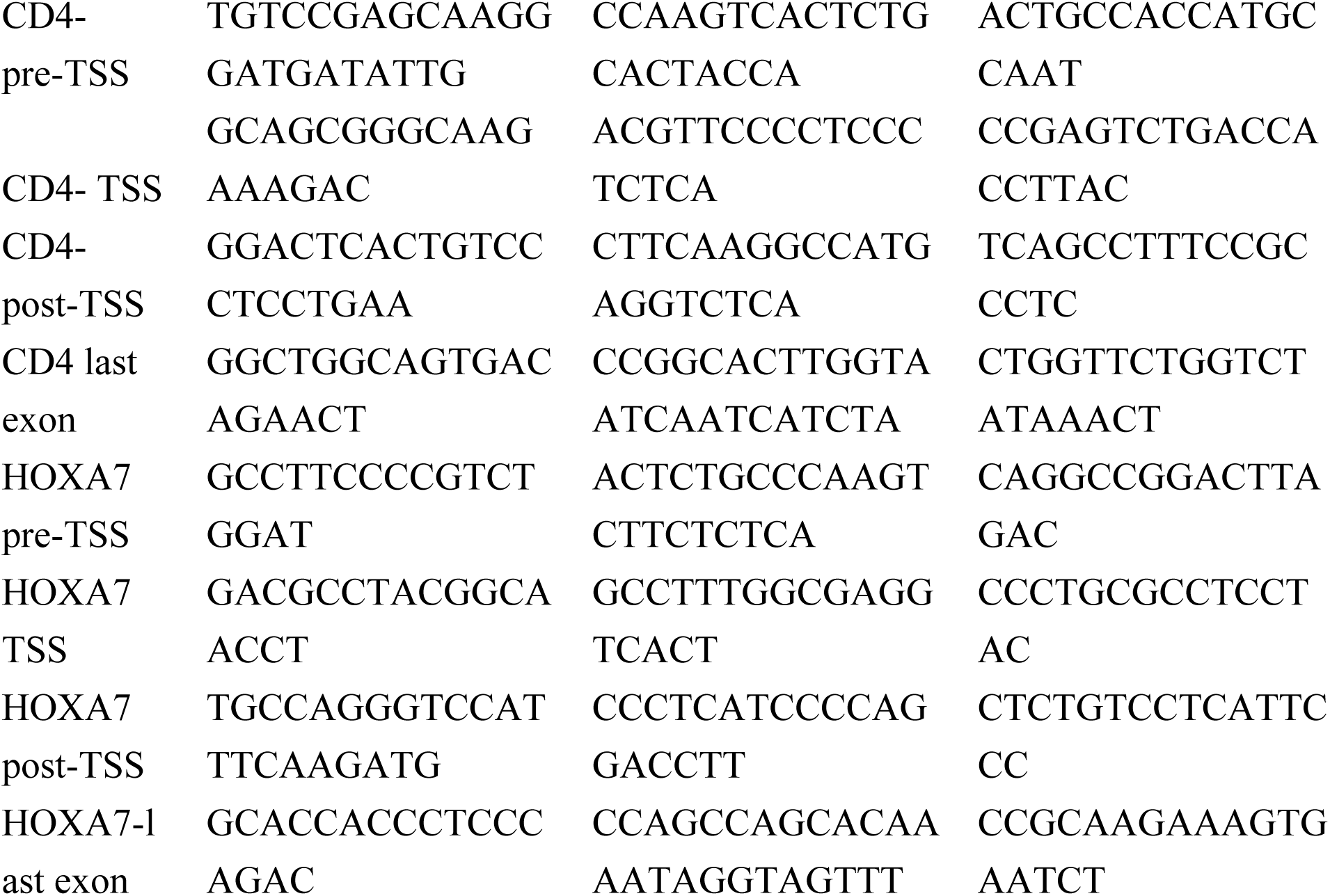
Custom qPCR probe/primer sequences.

**Table S4.**
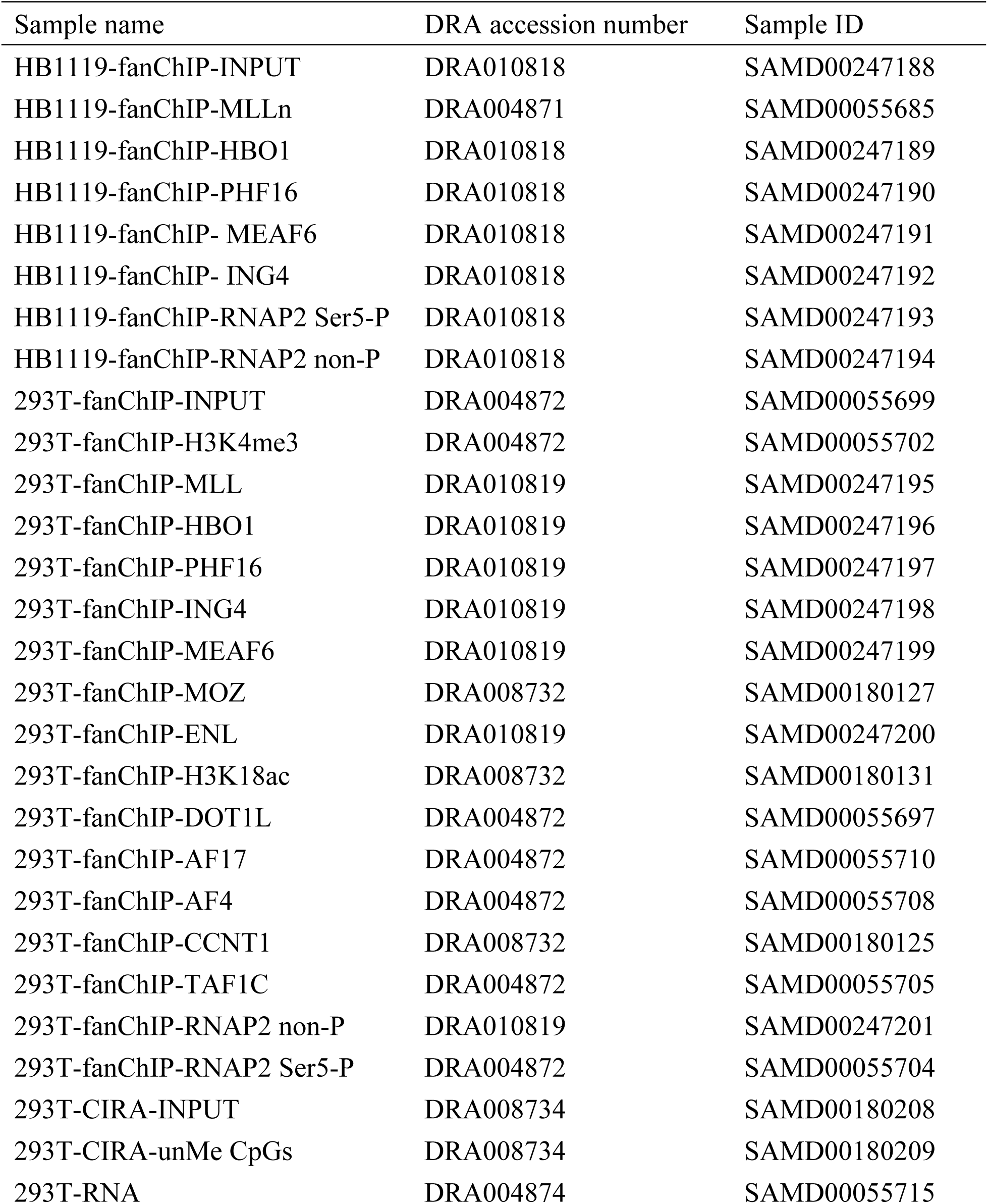
Accession numbers of NGS data.

| Sample name | DRA accession number | Sample ID |
| --- | --- | --- |
| HB1119-fanChIP-INPUT | DRA010818 | SAMD00247188 |
| HB1119-fanChIP-MLLn | DRA004871 | SAMD00055685 |
| HB1119-fanChIP-HBO1 | DRA010818 | SAMD00247189 |
| HB1119-fanChIP-PHF16 | DRA010818 | SAMD00247190 |
| HB1119-fanChIP- MEAF6 | DRA010818 | SAMD00247191 |
| HB1119-fanChIP- ING4 | DRA010818 | SAMD00247192 |
| HB1119-fanChIP-RNAP2 Ser5-P | DRA010818 | SAMD00247193 |
| HB1119-fanChIP-RNAP2 non-P | DRA010818 | SAMD00247194 |
| 293T-fanChIP-INPUT | DRA004872 | SAMD00055699 |
| 293T-fanChIP-H3K4me3 | DRA004872 | SAMD00055702 |
| 293T-fanChIP-MLL | DRA010819 | SAMD00247195 |
| 293T-fanChIP-HBO1 | DRA010819 | SAMD00247196 |
| 293T-fanChIP-PHF16 | DRA010819 | SAMD00247197 |
| 293T-fanChIP-ING4 | DRA010819 | SAMD00247198 |
| 293T-fanChIP-MEAF6 | DRA010819 | SAMD00247199 |
| 293T-fanChIP-MOZ | DRA008732 | SAMD00180127 |
| 293T-fanChIP-ENL | DRA010819 | SAMD00247200 |
| 293T-fanChIP-H3K18ac | DRA008732 | SAMD00180131 |
| 293T-fanChIP-DOT1L | DRA004872 | SAMD00055697 |
| 293T-fanChIP-AF17 | DRA004872 | SAMD00055710 |
| 293T-fanChIP-AF4 | DRA004872 | SAMD00055708 |
| 293T-fanChIP-CCNT1 | DRA008732 | SAMD00180125 |
| 293T-fanChIP-TAF1C | DRA004872 | SAMD00055705 |
| 293T-fanChIP-RNAP2 non-P | DRA010819 | SAMD00247201 |
| 293T-fanChIP-RNAP2 Ser5-P | DRA004872 | SAMD00055704 |
| 293T-CIRA-INPUT | DRA008734 | SAMD00180208 |
| 293T-CIRA-unMe CpGs | DRA008734 | SAMD00180209 |
| 293T-RNA | DRA004874 | SAMD00055715 |

## Additional information regarding the NGS data

ChIP-seq data, CIRA-seq data, and mRNA-seq data are deposited in DDBJ data archive and are available under the accession numbers and temporary access keys described below.

ChIP-seq and CIRA-seq files related to this study have been deposited in the DNA Data Bank of Japan (DDBJ) under the DRA accession numbers (GEA accession numbers) of DRA004871(E-GEAD-319), DRA004872(E-GEAD-320), DRA008732(E-GEAD-322), DRA008734(E-GEAD-324), DRA010818(E-GEAD-401), DRA010819(E-GEAD-402). The Fastq files of DRA004871(E-GEAD-319), DRA004872(E-GEAD-320),

total number of reads and uniquely mapped reads length of reads type of reads

HB1119-fanChIP-INPUT 37,157,634 32,242,893 50 single-end

HB1119-fanChIP-MLLn 27,823,993 22,362,095 50 single-end

HB1119-fanChIP-HBO1 48,063,278 41,419,128 50 single-end

HB1119-fanChIP-PHF16 39,286,468 33,831,669 50 single-end

HB1119-fanChIP-MEAF6 40,233,677 34,248,018 50 single-end

HB1119-fanChIP-ING4 37,184,884 31,018,894 50 single-end

HB1119-fanChIP-RNAP2 Ser5-P 37,064,277 32,941,328 50 single-end

HB1119-fanChIP-RNAP2 non-P 24,795,260 22,131,044 50 single-end

293T-fanChIP-INPUT 34,176,787 29,856,157 50 single-end

293T-fanChIP-H3K4me3 29,493,262 25,543,458 50 single-end

293T-fanChIP-MLL 30,988,174 26,294,921 50 single-end

293T-fanChIP-HBO1 29,580,779 25,601,332 50 single-end

293T-fanChIP-PHF16 27,550,541 23,664,906 50 single-end

293T-fanChIP-ING4 26,958,615 23,123,362 50 single-end

293T-fanChIP-MEAF6 27,884,294 24,082,140 50 single-end

293T-fanChIP-MOZ 27,874,246 20,175,414 50 single-end

293T-fanChIP-ENL 28,833,149 24,650,292 50 single-end

293T-fanChIP-H3K18ac 46,286,322 40,319,953 50 single-end

293T-fanChIP-DOT1L 26,657,236 22,155,835 50 single-end

293T-fanChIP-AF17 30,323,037 25,112,836 50 single-end

293T-fanChIP-AF4 29,290,866 22,783,533 50 single-end

293T-fanChIP-CCNT1 29,590,957 25,037,499 50 single-end

293T-fanChIP-TAF1C 29,729,743 24,939,481 50 single-end

293T-fanChIP-RNAP2 non-P 38,787,123 33,731,622 50 single-end

293T-fanChIP-RNAP2 Ser5-P 29,575,826 26,321,315 50 single-end

293T-CIRA-INPUT 22,076,417 18,287,840 50 single-end

293T-CIRA-unMe CpGs 22,131,576 10,799,441 50 single-end

## References

Artinger EL, Mishra BP, Zaffuto KM, Li BE, Chung EK, Moore AW, Chen Y, Cheng C, Ernst P (2013) An MLL-dependent network sustains hematopoiesis. Proc Natl Acad Sci U S A 110: 12000–12005

Au YZ, Gu M, De Braekeleer E, Gozdecka M, Aspris D, Tarumoto Y, Cooper J, Yu J, Ong SH, Chen X et al (2020) KAT7 is a genetic vulnerability of acute myeloid leukemias driven by MLL rearrangements. *Leukemia*

Avvakumov N, Lalonde ME, Saksouk N, Paquet E, Glass KC, Landry AJ, Doyon Y, Cayrou C, Robitaille GA, Richard DE et al (2012) Conserved molecular interactions within the HBO1 acetyltransferase complexes regulate cell proliferation. Mol Cell Biol 32: 689–703

DiMartino JF, Miller T, Ayton PM, Landewe T, Hess JL, Cleary ML, Shilatifard A (2000) A carboxy-terminal domain of ELL is required and sufficient for immortalization of myeloid progenitors by MLL-ELL. Blood 96: 3887–3893

Dull T, Zufferey R, Kelly M, Mandel RJ, Nguyen M, Trono D, Naldini L (1998) A third-generation lentivirus vector with a conditional packaging system. J Virol 72: 8463–8471

Erb MA, Scott TG, Li BE, Xie H, Paulk J, Seo HS, Souza A, Roberts JM, Dastjerdi S, Buckley DL et al (2017) Transcription control by the ENL YEATS domain in acute leukaemia. Nature 543: 270–274

Foy RL, Song IY, Chitalia VC, Cohen HT, Saksouk N, Cayrou C, Vaziri C, Cote J, Panchenko MV (2008) Role of Jade-1 in the histone acetyltransferase (HAT) HBO1 complex. J Biol Chem 283: 28817–28826

Goodfellow SJ, Zomerdijk JC (2013) Basic mechanisms in RNA polymerase I transcription of the ribosomal RNA genes. Sub-cellular biochemistry 61: 211–236

Gough SM, Slape CI, Aplan PD (2011) NUP98 gene fusions and hematopoietic malignancies: common themes and new biologic insights. Blood 118: 6247–6257

Hayashi Y, Harada Y, Kagiyama Y, Nishikawa S, Ding Y, Imagawa J, Shingai N, Kato N, Kitaura J, Hokaiwado S et al (2019) NUP98-HBO1-fusion generates phenotypically and genetically relevant chronic myelomonocytic leukemia pathogenesis. Blood Adv 3: 1047–1060

Heckl D, Kowalczyk MS, Yudovich D, Belizaire R, Puram RV, McConkey ME, Thielke A, Aster JC, Regev A, Ebert BL (2014) Generation of mouse models of myeloid malignancy with combinatorial genetic lesions using CRISPR-Cas9 genome editing. Nat Biotechnol 32: 941–946

Kaltenbach S, Soler G, Barin C, Gervais C, Bernard OA, Penard-Lacronique V, Romana SP (2010) NUP98-MLL fusion in human acute myeloblastic leukemia. Blood 116: 2332–2335

Klossowski S, Miao H, Kempinska K, Wu T, Purohit T, Kim E, Linhares BM, Chen D, Jih G, Perkey E et al (2019) Menin inhibitor MI-3454 induces remission in MLL1-rearranged and NPM1-mutated models of leukemia. *J Clin Invest*

Krivtsov AV, Evans K, Gadrey JY, Eschle BK, Hatton C, Uckelmann HJ, Ross KN, Perner F, Olsen SN, Pritchard T et al (2019) A Menin-MLL Inhibitor Induces Specific Chromatin Changes and Eradicates Disease in Models of MLL-Rearranged Leukemia. Cancer cell 36: 660–673 e611

Krivtsov AV, Twomey D, Feng Z, Stubbs MC, Wang Y, Faber J, Levine JE, Wang J, Hahn WC, Gilliland DG et al (2006) Transformation from committed progenitor to leukaemia stem cell initiated by MLL-AF9. Nature 442: 818–822

Kroon E, Thorsteinsdottir U, Mayotte N, Nakamura T, Sauvageau G (2001) NUP98-HOXA9 expression in hemopoietic stem cells induces chronic and acute myeloid leukemias in mice. EMBO J 20: 350–361

Lavau C, Szilvassy SJ, Slany R, Cleary ML (1997) Immortalization and leukemic transformation of a myelomonocytic precursor by retrovirally transduced HRX-ENL. Embo J 16: 4226–4237

Li H, Durbin R (2009) Fast and accurate short read alignment with Burrows-Wheeler transform. Bioinformatics 25: 1754–1760

Li Y, Wen H, Xi Y, Tanaka K, Wang H, Peng D, Ren Y, Jin Q, Dent SY, Li W et al (2014) AF9 YEATS domain links histone acetylation to DOT1L-mediated H3K79 methylation. Cell 159: 558–571

Liedtke M, Ayton PM, Somervaille TC, Smith KS, Cleary ML (2010) Self-association mediated by the Ras association 1 domain of AF6 activates the oncogenic potential of MLL-AF6. Blood 116: 63–70

Lin C, Smith ER, Takahashi H, Lai KC, Martin-Brown S, Florens L, Washburn MP, Conaway JW, Conaway RC, Shilatifard A (2010) AFF4, a component of the ELL/P-TEFb elongation complex and a shared subunit of MLL chimeras, can link transcription elongation to leukemia. Mol Cell 37: 429–437

Luo RT, Lavau C, Du C, Simone F, Polak PE, Kawamata S, Thirman MJ (2001) The elongation domain of ELL is dispensable but its ELL-associated factor 1 interaction domain is essential for MLL-ELL-induced leukemogenesis. Mol Cell Biol 21: 5678–5687

MacPherson L, Anokye J, Yeung MM, Lam EYN, Chan YC, Weng CF, Yeh P, Knezevic K, Butler MS, Hoegl A et al (2020) HBO1 is required for the maintenance of leukaemia stem cells. Nature 577: 266–270

Martin M (2011) Cutadapt Removes Adapter Sequences From High-Throughput Sequencing Reads. EMBnet Journal 17: 10–12

Meyer C, Burmeister T, Groger D, Tsaur G, Fechina L, Renneville A, Sutton R, Venn NC, Emerenciano M, Pombo-de-Oliveira MS et al (2018) The MLL recombinome of acute leukemias in 2017. Leukemia 32: 273–284

Milne TA, Kim J, Wang GG, Stadler SC, Basrur V, Whitcomb SJ, Wang Z, Ruthenburg AJ, Elenitoba-Johnson KS, Roeder RG et al (2010) Multiple interactions recruit MLL1 and MLL1 fusion proteins to the HOXA9 locus in leukemogenesis. Mol Cell 38: 853–863

Mishima Y, Miyagi S, Saraya A, Negishi M, Endoh M, Endo TA, Toyoda T, Shinga J, Katsumoto T, Chiba T et al (2011) The Hbo1-Brd1/Brpf2 complex is responsible for global acetylation of H3K14 and required for fetal liver erythropoiesis. Blood 118: 2443–2453

Miyamoto R, Okuda H, Kanai A, Takahashi S, Kawamura T, Matsui H, Kitamura T, Kitabayashi I, Inaba T, Yokoyama A (2020) Activation of CpG-Rich Promoters Mediated by MLL Drives MOZ-Rearranged Leukemia. Cell Rep 32: 108200

Morita S, Kojima T, Kitamura T (2000) Plat-E: an efficient and stable system for transient packaging of retroviruses. Gene Ther 7: 1063–1066

Nagase T, Yamakawa H, Tadokoro S, Nakajima D, Inoue S, Yamaguchi K, Itokawa Y, Kikuno RF, Koga H, Ohara O (2008) Exploration of human ORFeome: high-throughput preparation of ORF clones and efficient characterization of their protein products. DNA Res 15: 137–149

Okuda H, Kanai A, Ito S, Matsui H, Yokoyama A (2015) AF4 uses the SL1 components of RNAP1 machinery to initiate MLL fusion- and AEP-dependent transcription. Nat Commun 6: 8869

Okuda H, Kawaguchi M, Kanai A, Matsui H, Kawamura T, Inaba T, Kitabayashi I, Yokoyama A (2014) MLL fusion proteins link transcriptional coactivators to previously active CpG-rich promoters. Nucleic Acids Res 42: 4241–4256

Okuda H, Stanojevic B, Kanai A, Kawamura T, Takahashi S, Matsui H, Takaori-Kondo A, Yokoyama A (2017) Cooperative gene activation by AF4 and DOT1L drives MLL-rearranged leukemia. J Clin Invest 127: 1918–1931

Okuda H, Takahashi S, Takaori-Kondo A, Yokoyama A (2016) TBP loading by AF4 through SL1 is the major rate-limiting step in MLL fusion-dependent transcription. Cell cycle 15: 2712–2722

Okuda H, Yokoyama A (2017a) In vivo Leukemogenesis Model Using Retrovirus Transduction. Bio-protocol 7

Okuda H, Yokoyama A (2017b) Myeloid Progenitor Transformation Assay. Bio-protocol 7

Qi S, Li Z, Schulze-Gahmen U, Stjepanovic G, Zhou Q, Hurley JH (2017) Structural basis for ELL2 and AFF4 activation of HIV-1 proviral transcription. Nat Commun 8: 14076

Shi A, Murai MJ, He S, Lund G, Hartley T, Purohit T, Reddy G, Chruszcz M, Grembecka J, Cierpicki T (2012) Structural insights into inhibition of the bivalent menin-MLL interaction by small molecules in leukemia. Blood 120: 4461–4469

Shilatifard A, Lane WS, Jackson KW, Conaway RC, Conaway JW (1996) An RNA polymerase II elongation factor encoded by the human ELL gene. Science 271: 1873–1876

Shima Y, Yumoto M, Katsumoto T, Kitabayashi I (2017) MLL is essential for NUP98-HOXA9-induced leukemia. Leukemia 31: 2200–2210

Simone F, Luo RT, Polak PE, Kaberlein JJ, Thirman MJ (2003) ELL-associated factor 2 (EAF2), a functional homolog of EAF1 with alternative ELL binding properties. Blood 101: 2355–2362

Simone F, Polak PE, Kaberlein JJ, Luo RT, Levitan DA, Thirman MJ (2001) EAF1, a novel ELL-associated factor that is delocalized by expression of the MLL-ELL fusion protein. Blood 98: 201–209

Takahashi H, Parmely TJ, Sato S, Tomomori-Sato C, Banks CA, Kong SE, Szutorisz H, Swanson SK, Martin-Brown S, Washburn MP et al (2011) Human mediator subunit MED26 functions as a docking site for transcription elongation factors. Cell 146: 92–104

Takahashi S, Yokoyama A (2020) The molecular functions of common and atypical MLL fusion protein complexes. Biochim Biophys Acta Gene Regul Mech 1863: 194548

Tkachuk DC, Kohler S, Cleary ML (1992) Involvement of a homolog of Drosophila trithorax by 11q23 chromosomal translocations in acute leukemias. Cell 71: 691–700

Tzelepis K, Koike-Yusa H, De Braekeleer E, Li Y, Metzakopian E, Dovey OM, Mupo A, Grinkevich V, Li M, Mazan M et al (2016) A CRISPR Dropout Screen Identifies Genetic Vulnerabilities and Therapeutic Targets in Acute Myeloid Leukemia. Cell Rep 17: 1193–1205

Wan L, Wen H, Li Y, Lyu J, Xi Y, Hoshii T, Joseph JK, Wang X, Loh YE, Erb MA et al (2017) ENL links histone acetylation to oncogenic gene expression in acute myeloid leukaemia. Nature 543: 265–269

Wang GG, Cai L, Pasillas MP, Kamps MP (2007) NUP98-NSD1 links H3K36 methylation to Hox-A gene activation and leukaemogenesis. Nat Cell Biol 9: 804–812

Wang Z, Song J, Milne TA, Wang GG, Li H, Allis CD, Patel DJ (2010) Pro isomerization in MLL1 PHD3-bromo cassette connects H3K4me readout to CyP33 and HDAC-mediated repression. Cell 141: 1183–1194

Wiederschain D, Kawai H, Gu J, Shilatifard A, Yuan ZM (2003) Molecular basis of p53 functional inactivation by the leukemic protein MLL-ELL. Mol Cell Biol 23: 4230–4246

Xu H, Valerio DG, Eisold ME, Sinha A, Koche RP, Hu W, Chen CW, Chu SH, Brien GL, Park CY et al (2016) NUP98 Fusion Proteins Interact with the NSL and MLL1 Complexes to Drive Leukemogenesis. Cancer cell 30: 863–878

Yokoyama A, Cleary ML (2008) Menin critically links MLL proteins with LEDGF on cancer-associated target genes. Cancer cell 14: 36–46

Yokoyama A, Kitabayashi I, Ayton PM, Cleary ML, Ohki M (2002) Leukemia proto-oncoprotein MLL is proteolytically processed into 2 fragments with opposite transcriptional properties. Blood 100: 3710–3718

Yokoyama A, Lin M, Naresh A, Kitabayashi I, Cleary ML (2010) A higher-order complex containing AF4 and ENL family proteins with P-TEFb facilitates oncogenic and physiologic MLL-dependent transcription. Cancer cell 17: 198–212

Yokoyama A, Somervaille TC, Smith KS, Rozenblatt-Rosen O, Meyerson M, Cleary ML (2005) The menin tumor suppressor protein is an essential oncogenic cofactor for MLL-associated leukemogenesis. Cell 123: 207–218

Yokoyama A, Wang Z, Wysocka J, Sanyal M, Aufiero DJ, Kitabayashi I, Herr W, Cleary ML (2004) Leukemia proto-oncoprotein MLL forms a SET1-like histone methyltransferase complex with menin to regulate Hox gene expression. Mol Cell Biol 24: 5639–5649

Yu BD, Hanson RD, Hess JL, Horning SE, Korsmeyer SJ (1998) MLL, a mammalian trithorax-group gene, functions as a transcriptional maintenance factor in morphogenesis. Proc Natl Acad Sci U S A 95: 10632–10636

Zhang Y, Guo Y, Gough SM, Zhang J, Vann KR, Li K, Cai L, Shi X, Aplan PD, Wang GG et al (2020) Mechanistic insights into chromatin targeting by leukemic NUP98-PHF23 fusion. Nat Commun 11: 3339

## References

Stewart SA, Dykxhoorn DM, Palliser D, Mizuno H, Yu EY, An DS, Sabatini DM, Chen IS, Hahn WC, Sharp PA et al (2003) Lentivirus-delivered stable gene silencing by RNAi in primary cells. RNA 9: 493–501

Thorvaldsdottir H, Robinson JT, Mesirov JP (2013) Integrative Genomics Viewer (IGV): high-performance genomics data visualization and exploration. Brief Bioinform 14: 178–192

